# Whole-brain receptor markers and hub excitabilities determine acute and long-term outcomes in stroke

**DOI:** 10.64898/2026.10.04.755271

**Authors:** Julian Schulte, Jakub Vohryzek, Martí Monge i Asensio, Gorka Zamora-López, Gustavo Patow, Yonatan Sanz Perl, Maurizio Corbetta, Gustavo Deco

## Abstract

Following a stroke, changes in receptor systems have been observed that may influence the excitability of affected brain regions and drive recovery trajectories. Translating results from animal studies to patients remains a major challenge, and while imaging modalities have provided a wealth of findings, a mechanistic link between local receptor systems and observed imaging data is still missing. To bridge this gap, we built patient-specific digital twins with their own neuromodulatory profiles: we integrated 19 receptor and transporter density maps into a mechanistic whole-brain model, and estimated their influence on cortical excitability for each stroke patient and how these systems subsequently explain recovery trajectories. We found that distinct receptor profiles modulated excitability in perilesional versus non-perilesional tissue. Favourable acute motor outcomes were most strongly explained by increased inhibition mediated by non-perilesional GABA-A receptors, together with enhanced excitability driven by cannabinoid, histaminergic, and noradrenergic systems. For acute language outcomes, we identified receptor signatures that, together with motor findings, formed two cross-domain clusters linking excitability to either better or poorer performance. Remarkably, structurally highly connected hubs played a key role: their acute excitability explained cognitive and language deficits and predicted language recovery at one year. Together, our results establish a mechanistic link between receptor-specific modulation of cortical excitability and behavioural recovery, highlighting potential targets for pharmacological and non-invasive therapeutic interventions.

## Introduction

Stroke frequently leads to lasting motor and cognitive disabilities, globally placing a substantial burden on those affected and their support networks. Elucidating the biological mechanisms that shape acute and long-term behavioural outcomes is therefore essential. At the neuronal level, a range of receptor and transporter alterations have been identified, some potentially promoting and others impairing acute function and later recovery. Among these, several alterations have been observed across neuromodulatory and the major excitatory and inhibitory receptor systems.

Early and subsequent animal work has highlighted GABAergic changes, showing disinhibition in perilesional and remote cortex, reflected by decreases in GABA-A markers (Alia et al., 2016; Neumann-Haefelin et al., 1995; Que, Witte, et al., 1999; Schiene et al., 1996). While higher initial GABA-A availability in patients across several brain regions predicted better short- and long-term motor outcomes (Kim et al., 2014), the same study also demonstrated that larger subsequent reductions in GABA-A availability, indicating disinhibition, were associated with greater motor improvement over time. The early protective role of GABA signalling may help mitigate glutamate-induced excitotoxicity (Lai et al., 2014; Ward, 2017). However, this balance appears to reverse over time as reduced inhibition and enhanced glutamatergic activity seem to become beneficial for recovery by potentially promoting neuroplasticity (Ward, 2017). Interestingly, an increase in bilateral NMDA receptor (NMDAR) availability (Que, Schiene, et al., 1999) and beneficial effects for cognitive and somatosensory outcomes after the intervention with an NMDAR partial agonist were observed in a stroke animal model (Dhawan et al., 2011). However, it remains unclear whether the observed benefits of the pharmacological intervention depend on the time-point after the stroke and/or the affected NMDAR subtypes and locations (Ge et al., 2020).

In addition to the main excitatory and inhibitory neurotransmitter systems, the involvement of multiple neuromodulatory systems in post-stroke pathophysiology and functional outcomes has been observed, for instance the cannabinoid (Bietar et al., 2023; Caltana et al., 2015; Carter et al., 2024), histaminergic (Hu & Chen, 2012; Lozada et al., 2005), serotonergic (Møller et al., 2007; Véra et al., 1996; Zahrai et al., 2020), and noradrenergic (Mizuki et al., 1995; L. E. Wang et al., 2011; Zahrai et al., 2020) systems, which may also directly or indirectly shape the excitability of brain regions after stroke. Interestingly, recent work has also identified distinct clusters of stroke-related neurotransmitter injury profiles that were associated with certain behavioural outcomes (Alves et al., 2025). Despite an abundance of findings, translating results from animal models to human patients remains a major challenge, and although positron emission tomography (PET) studies have provided invaluable insights into the receptor and transporter landscape of the brain (Hansen et al., 2022) and their post-stroke alterations, they do not offer direct measures of cortical excitability (Ward, 2017) or crucially mechanistic explanations.

To bridge this gap, the principal goal of this study was to investigate how a multitude of receptor and transporter systems shape neuronal cortical excitability and, in turn, explain post-stroke behavioural outcomes at the acute (1–2 weeks) and long-term (1 year) stages. Building on prior work incorporating heterogeneity information into whole-brain models (Deco et al., 2021, 2026; Demirtaş et al., 2019; Monge i Asensio, 2025; Sanz Perl, Zamora-Lopez, et al., 2023), we have built a digital twin for each patient with their own neuromodulatory profiles: From an independent atlas, we integrated a total of 19 cortical receptor and transporter distributions (Hansen et al., 2022) and estimated the influence of these general maps on cortical excitability for each patient by fitting the twins to the individual empirical imaging data. The influence of these maps was independently estimated for perilesional and non-perilesional regions, accounting for the distinct pathophysiology of the area surrounding the stroke (Murphy & Corbett, 2009). This allowed us to determine the influence of different receptor and transporter maps on cortical excitability for each patient and link these characteristics to behavioural outcomes.

We found that favourable acute motor outcomes were most strongly associated with increased inhibition mediated by non-perilesional GABA-A receptors and with an enhanced excitability driven by non-perilesional cannabinoid (CB1) as well as by perilesional histaminergic (H3) and norepinephrine systems. For acute language outcomes, we identified several significant receptors and transporters, a subset of which overlapped with the motor findings to form two cross-domain clusters where increased excitability was associated with either better or poorer outcomes across both functions. Investigating individual cortical brain regions revealed that higher interconnectivity and greater structural damage to white matter pathways were associated with reduced excitability. Importantly, the acute excitability of highly central brain regions (i.e., hubs) was decisive both for acute cognitive and language outcomes, and for predicting language recovery one year after the stroke. Interestingly, the contribution of excitability to these longitudinal language outcomes was modulated by the hub’s structural integrity. These results illustrate that our patient-specific whole-brain modelling framework can recover relevant receptors and transporters to explain acute deficits, while also identifying the excitability of hubs as a central driver of both short- and long-term functional outcomes.

## Methods

### Overview - Receptor-informed personalized whole-brain model as digital twin

The principal goal of this study was to assess how cortical receptor and transporter distributions, along with the regional excitabilities they shape, explain motor, language, and general cognitive outcomes in the acute (T1; 1-2 weeks) and long-term (T3; one year) stages after a stroke. To estimate their influence on cortical excitability, we employed a patient-specific whole-brain model i.e., digital twin fitted to clinical data. The clinical dataset comprised fMRI and behavioural measurements from 96 patients in the acute stage and 49 patients at follow-up (Fig. 1a). The neurochemical architecture informing these models was derived from an independent receptor atlas and comprised 19 cortical receptor and transporter density maps (Fig. 1b; see *Receptor density maps* for the full list). We integrated this biological information into our twins by linking regional receptor and transporter densities to neuronal excitability. Specifically, each brain region was modelled as coupled excitatory and inhibitory neuronal populations, and regional excitability was quantified based on the excitatory population, representing its firing sensitivity for a given synaptic input current (Fig. 1c). Regional excitability was then derived as a linear combination of the empirical density maps, weighted by model-derived receptor-specific parameters (Fig. 1d). These parameters, referred to as *receptor scalings*, captured the contribution of each map to cortical excitability and were estimated by fitting the whole-brain model to each patient’s empirical data (Fig. 1e). To account for lesion-related differences, we estimated the receptor scalings separately for perilesional and non-perilesional regions (Fig. 1d).

**Figure 1.**
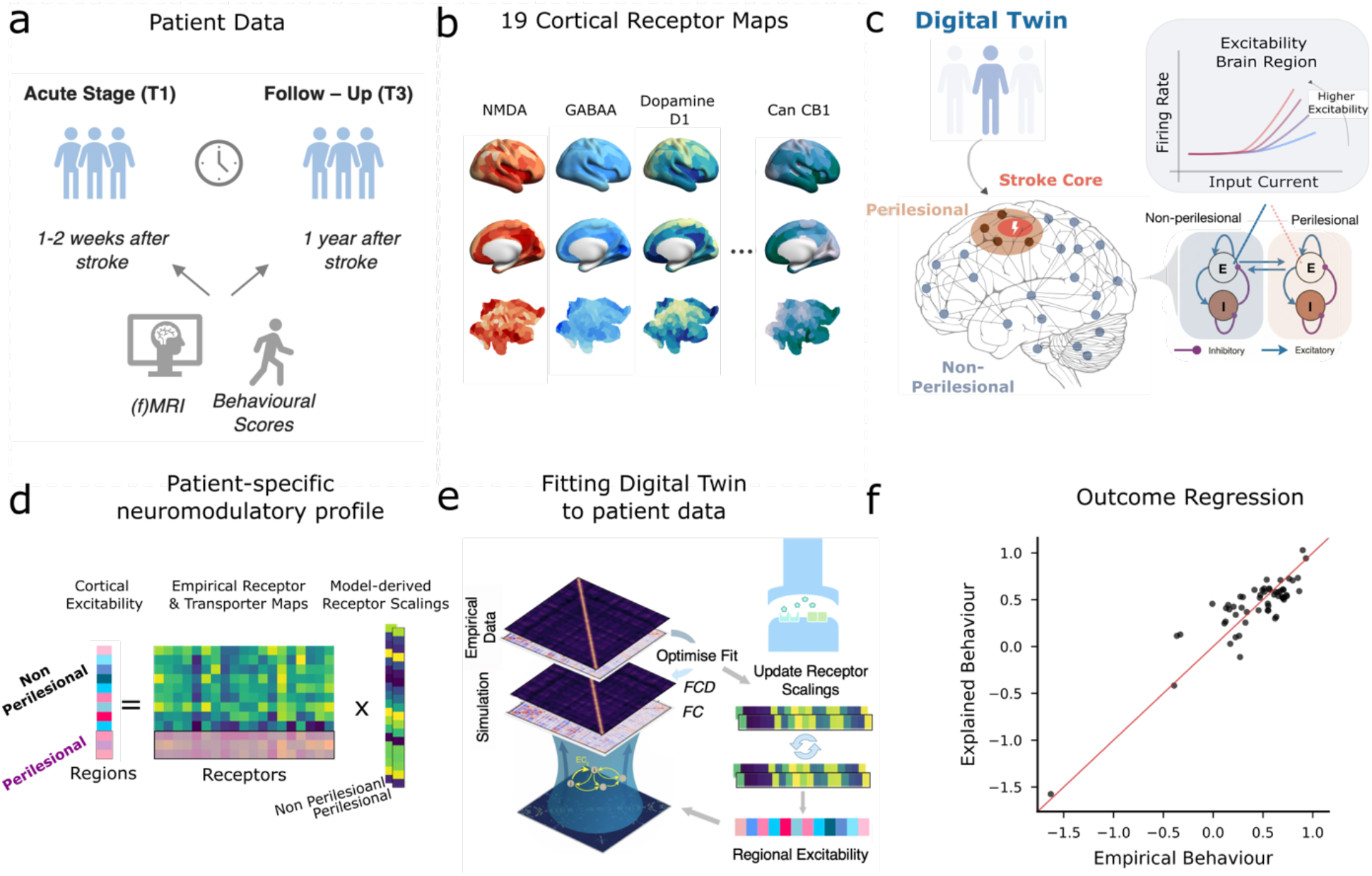
Receptor-informed whole-brain model as digital twin. **a**| Clinical (f)MRI and behavioural data were collected from stroke patients during the acute stage (T1; 1–2 weeks post-stroke) and at one-year follow-up (T3). **b**|To capture spatial receptor and transporter distributions, we utilised an independent atlas comprising 19 cortical maps. **c**|We incorporated these density maps in our whole-brain model i.e., digital twin to determine the neuronal excitability of all cortical brain regions. Each brain region was modelled as a coupled excitatory– inhibitory (E–I) circuit, with regional excitability defined as the gain of the excitatory population, that is, the sensitivity of its firing rate to input current. **d**|Specifically, regional neuronal excitability (left) was derived as a linear combination of the empirical receptor and transporter densities (middle) and patient-specific receptor-specific scaling parameters, which were derived from the model (right). These scaling parameters were estimated separately for perilesional and non-perilesional areas and reflect the influence of a receptor or transporter map on cortical excitability. **e**|The scaling parameters were obtained from the whole-brain model by fitting simulated functional connectivity (FC) and functional connectivity dynamics (FCD) to the empirical data. **f**|The optimized patient-specific receptor scalings and excitability values were subsequently tested for their ability to explain acute and predict long-term behavioural outcomes after stroke.

To fit the model, we estimated the receptor scaling parameters, a global coupling factor, and directly subcortical excitability values to reproduce each patient’s functional connectivity (FC) and functional connectivity dynamics (FCD; Fig. 1e). Brain regions in the model were interlinked according to an average healthy effective connectivity map (EC) masked with patient lesion information. The applied framework enabled the estimation of regional cortical excitabilities and the influence of individual receptor and transporter maps, formalised through the receptor scaling (“receptor scaling” refers to both receptor and transporter map scaling). The fitted scalings quantify each map’s contribution to neuronal excitability. The contribution of the empirical receptor or transporter maps on cortical excitability may reflect direct but also indirect underlying biological mechanisms, the latter being particularly likely for transporters. As a final step, we examined how the scaling parameters and the cortical excitability values derived from leading brain hubs related to motor, language, and cognitive outcomes in stroke patients (Fig. 1f).

### Subjects

A longitudinal dataset of stroke patients from the Washington University Stroke Cohort (Corbetta et al., 2015) was used. We examined data collected within 1-2 weeks (acute stage; T1) and 1 year (long-term stage; T3) post-stroke. Ischemic stroke accounted for the majority of cases (83%), with 17% of patients having suffered hemorrhagic strokes. Detailed data collection protocols are described in Corbetta et al. (2015). Patient data were acquired through the stroke service at Barnes-Jewish Hospital (BJH) together with the Washington University Cognitive Rehabilitation Research Group (CRRG), with recruitment focused on individuals experiencing a first-time, single-lesion stroke. A total of *N* = 96 stroke patients were included. To generate the healthy effective connectivity, we also included data from 27 age-matched healthy control participants. Neuroimaging data, encompassing both structural and functional measures, were acquired concurrently with neuropsychological assessments.

### Neuropsychological and behavioural data

For the acute and long-term stages, participants performed a broad neuropsychological testing battery. In total, 44 behavioural measures were obtained, covering motor performance, language abilities, attentional function, and memory processes (Corbetta et al., 2015). To obtain compact representations of performance within each domain, a Principal Component Analysis (PCA) was applied separately to the test scores of each functional category. This procedure yielded domain-specific summary scores, including MotorR and MotorL (right and left body motor performance), Language, MemoryS (spatial memory composite), MemoryV (verbal memory composite), AttentionVF (visuospatial field bias), mean attentional performance across tasks, and AttentionValDis (ability to reorient attention toward previously unattended stimuli). Motor performance was assessed using tasks probing both upper and lower extremity function. Behavioural scores were standardized relative to the healthy control group using z-transformation, enabling identification of domain-specific impairments (see Corbetta et al., 2015 for methodological details). Subsequently, our analysis focused on general motor, language, and cognitive outcomes. To derive global performance metrics, we averaged all functional motor subdomains into a general motor score and all memory and attention measures into a general cognition score. As behavioural data were not universally available across all time points, we specified the exact number of included participants for each separate analysis.

### Neuroimaging acquisition and pre-processing

Imaging was performed using a Siemens 3T Tim-Trio scanner (12-channel head coil) at Washington University School of Medicine. The acquisition protocol utilized a sagittal T1-weighted MP-RAGE sequence (TR = 1950 ms, TE = 2.26 ms, flip angle = 90°, 1.0 mm^3^ voxels) alongside gradient-echo EPI (TR = 2000 ms, TE = 2 ms, 4 × 4 mm in-plane resolution, 32 slices). Participants completed between six and eight resting-state runs (∼30 minutes total; 896 time points), during which they were directed to fixate on a white cross displayed on a black background (Griffis et al., 2019). Resting-state fMRI data underwent a standard pre-processing pipeline. This involved the regression of nuisance signals (white matter, cerebrospinal fluid, and ventricles), correction for head motion, and temporal filtering (0.008–0.08 Hz). Frames exceeding a framewise displacement (FD) threshold of 0.5 mm were removed. The cleaned BOLD time series were mapped onto a set of 234 regions of interest (ROIs). The Schaefer atlas (Schaefer et al., 2018) was utilized for the 200 cortical parcels, the AAL (Tzourio-Mazoyer et al., 2002) for the 34 subcortical and cerebellar regions, and the Harvard–Oxford Subcortical atlas for brainstem identification. Finally, a detrending of the BOLD signal was performed.

A structural connectome atlas was generated from diffusion MRI streamline tractography using high-angular Human Connectome Project (HCP) data from 842 healthy participants (Yeh et al., 2018). In MNI template space, this atlas (Griffis et al., 2019, 2021) leverages high-spatial and angular resolution diffusion imaging together with q-space diffeomorphic reconstruction (Yeh & Tseng, 2011). By averaging spin distribution functions across the population and applying deterministic tractography with multiple turning-angle thresholds, 500,000 streamline trajectories were generated. The resulting group-averaged connectivity matrix defines the anatomical edge weights between the study’s brain regions.

### Stroke Lesions & disconnection masks

Structural MRI images were manually segmented using the biomedical imaging software system Analyze (Robb & Hanson, 1991) and subsequently classified with an unsupervised K-means clustering algorithm that determined the overlap with white matter, gray matter, and subcortical masks (including the thalamus and basal ganglia; see Corbetta et al., 2015). Consistent with previous studies (Idesis et al., 2024; Sarasso et al., 2020), we designated the 1 cm zone (Euclidean distance) immediately surrounding the lesion as the perilesional area. The initial cohort consisted of 96 subjects in the acute stage. Six patients were removed from analyses requiring perilesional mapping because this region could not be reliably defined within the 1 cm threshold. Structural connectivity damage was evaluated using the Lesion Quantification Toolkit (Griffis et al., 2021). By referencing population-level atlases of grey matter parcels and white matter trajectories (Yeh et al., 2018), this tool calculates specific damage estimates. Consequently, a patient-specific Structural Disconnection (SDC) mask was generated for each subject, yielding a whole-brain matrix that quantifies the extent of remaining intact connections between every pair of regions.

### Regional degree and structural connectivity damage

To identify the most structurally connected brain regions, we computed regional degree from the binarized average healthy structural connectome, assigning existing connections a value of one and absent connections a value of zero. Degree was defined as the total number of connections for each brain region. To account for patient-specific structural disconnections in our statistical analyses, we calculated a regional damage score for each individual. Specifically, the average healthy structural connectivity matrix was element-wise multiplied by the SDC mask of a patient, and the loss in connection strength was summed across all edges connected to a given region. Repeating this procedure for all patients generated a subject-by-region matrix quantifying the magnitude of structural connectivity impairment for every brain region in each patient. To reduce the dimensionality, we applied Principal Component Analysis (PCA), retaining the leading components that explained approximately 65% of the variance in the behavioural outcome regressions (see Behavioural outcome regressions).

### Receptor density maps

To enrich our whole-brain modelling framework with biological information, we included multiple receptor and transporter density markers, which could potentially influence the excitability of cortical brain regions after stroke. This approach builds on previous modelling work that incorporated regional heterogeneity information (Deco et al., 2021, 2026; Demirtaş et al., 2019; Monge i Asensio, 2025; Sanz Perl, Zamora-Lopez, et al., 2023), extending whole-brain network models to include receptor-level information in a patient-specific stroke context. We used the publicly accessible dataset by Hansen et al. (2022), which comprises nine neurotransmitter systems obtained from over 1200 healthy participants. In detail, we included the cortical receptor and transporter estimation of dopamine (D1, D2, DAT), norepinephrine (NET), serotonin (5HT1A, 5HT1B, 5HT2A, 5HT4, 5HT6, 5HTT), acetylcholine (A4B2, M1, VAChT), glutamate (mGLUR5, NMDA), GABA (GABA-A), histamine (H3), cannabinoid (CB1), and opioid (MOR). The positron emission tomography (PET) data were spatially registered to the MNI-ICBM 152 nonlinear 2009 template (version c, asymmetric), then averaged for each group of participants within the respective research projects, before being parcellated. If more than one dataset was available for a single receptor or transporter, we selected the dataset with the largest number of subjects. As a final step, we applied min–max normalization to the cortical density maps.

### Surrogate receptor maps

To check if our whole-brain modelling results were driven by the spatial topography of the biological receptor maps, we constructed spatially autocorrelated surrogate maps as a null-model for each of the 19 receptor and transporter type. For this, we used the Moran spectral randomization procedure (Wagner & Dray, 2015) supplied in the Neuromaps library (Markello et al., 2022). For each patient, we generated a new set of reshuffled surrogate maps for each receptor and transport types. As with the biological maps, we applied in a last step the min– max normalization. Whole-brain models were subsequently re-fitted using surrogate maps, and the resulting model error was compared against that obtained with the biological receptor maps.

### Estimation of effective connectivity

Generative Effective Connectivity (GEC) is a model-based quantification of the directional influence between brain regions. Previous work has demonstrated that combining this metric with data on structural dysconnectivity supports the simulation of patient-specific functional connectivity (Idesis et al., 2024). Utilizing this framework, we derived the GEC to serve as the structural scaffold together with the SDC maps for our whole-brain simulations. Specifically, we first estimated the GEC from a healthy reference cohort (*N* = 27) following the optimization protocol of Deco et al. (2019). We then masked this healthy GEC with patient-specific SDC maps (via element-wise multiplication), explicitly embedding the individual structural disconnections. For the GEC calculation itself, BOLD time series were simulated using a Hopf model. In this architecture, local dynamics were defined by the normal form of a supercritical Hopf bifurcation, with inter-regional coupling governed by the empirical structural connectivity matrix *C*. The activity of each region is given by:

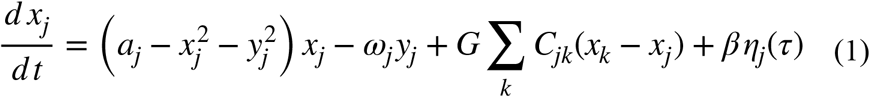

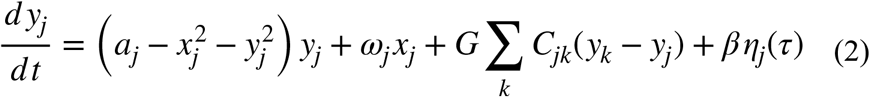

The real component *x*_*j*_ (1) and the imaginary component *y*_*j*_ (2) define the system’s dynamics. Following standard protocols, we fixed the bifurcation parameter near the Hopf bifurcation i.e., *a* (Deco et al., 2017). Intrinsic nodal frequencies (ω) were estimated using the mean peak frequency of the narrowband empirical BOLD signals. The interaction between regions is modulated by a global coupling strength and Gaussian noise (η) is also added with a standard deviation of *β* = 0.04. Finally, we applied a scaling factor of 0.2 to the structural connectivity weights to normalize them and optimized the model by minimizing the phase mismatch between the simulated and healthy empirical functional connectivity (at).

Once the whole-brain Hopf model was optimized, we derived the GEC by sequentially updating the structural connectivity weights (*C*) using a gradient descent strategy. This approach minimized the distance between the empirical and model-derived grand-average phase coherence matrices. We continued this procedure until the fit stabilized, consistent with prior work (Deco et al., 2023; Idesis et al., 2022; Sanz Perl, Geli, et al., 2023). In detail, connectivity weights were adjusted according to:

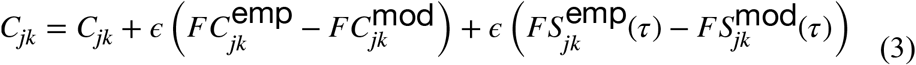

The deviations between the average empirical and model functional phase coherence matrices 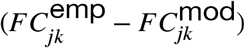, combined with the time-shifted covariance differences 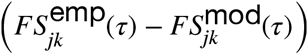, were weighted by *ϵ* = 0.001 and added to the preceding connections. We calculated the covariance matrices with a time-shift of *τ*, normalizing them across node pairs by 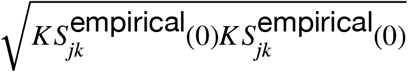. Comprehensive methodological details can be found in Idesis et al. (2024).

### Whole-brain simulation – Biophysical Dynamic Mean Field model

Patient-specific whole-brain simulations were conducted using the Dynamic Mean Field (DMF) model, originally introduced by Deco et al. (2014). This biophysical framework models spontaneous neural activity by simulating the interactions of localized neural populations (ROIs) connected via anatomical pathways. Building on the seminal work of Wong and Wang (2006), the model reduces the complex dynamics of large networks of excitatory and inhibitory integrate-and-fire spiking neurons to a simplified system of equations describing the interactions between coupled excitatory (E) and inhibitory (I) populations. Locally, each brain region comprises an E-I population pair, with dynamics driven by both internal recurrent signalling and mutual interactions. Synaptic activity is modelled, where NMDA/ AMPA receptors mediate excitatory currents and GABA-A receptors govern inhibitory currents. Long-distance communication is transmitted exclusively through excitatory projections, weighted by the structural connectivity *C* and modulated by a global scaling parameter . Crucially, for these simulations, *C* was defined as the GEC masked by the patient’s specific structural disconnection (SDC; as outlined before in Generative Effective Connectivity). The activity within a single brain region is described by:

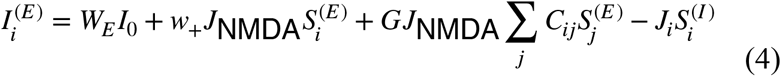

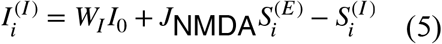

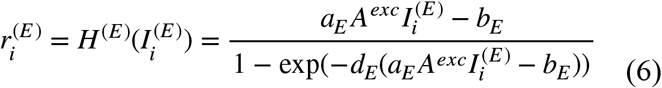

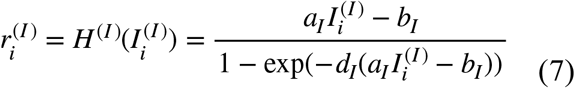

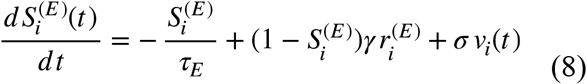

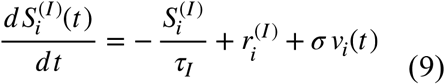

Within the model equations, 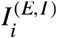, 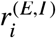, and 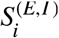 represent the input current, firing rate, and synaptic gating, respectively for the excitatory or inhibitory populations. The external input current *I*_0_ = 0.382(*n A*) is weighted by population-specific scaling factors *W*_*E*_ = 1 or *W*_*I*_ = 0.7. The excitatory synaptic coupling is denoted by *J*_*NMDA*_ = 0.15(*n A*) and *w*_+_ = 1.4 signifies local recurrent excitation within the same region. To suppress excessive excitation, *J*_*i*_ represents the local inhibitory feedback. This feedback parameter is calculated using the efficient algorithm proposed by Herzog et al. (2024), defined as 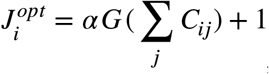, with *α* = 0.75.

*H* ^(*E*/*I* )^ denotes the transfer function and maps synaptic input currents to population firing rates. Importantly, to parametrize the excitability of the excitatory population, the local excitatory current *I* ^(*E* )^ is scaled by *A*^*exc*^ (Equation 6). This parameter captures the firing sensitivity of excitatory neurons to synaptic inputs, thereby encapsulating population-level excitability. Specifically, *A*^*exc*^ is a vector representing the excitability for each brain region. We derive it for cortical regions by estimating the additive influence of 19 different receptor and transporter density maps on this excitability (detailed in *Introducing receptor map heterogeneity*). For subcortical brain regions, we directly estimate it without receptor map information. All other parameters in the input-output functions are held constant and detailed in Table 1. Kinetic parameters are set as *τ*_*E*_ = *τ*_*NMDA*_ = 100*ms, τ*_*I*_ = *τ*_*GABA*_ = 10*ms*, and *γ* = 0.641/1000 (with a factor of 1,000 to express time constants in milliseconds). Uncorrelated Gaussian noise *v*_*i*_(*t* ) with standard deviation *σ* = 0.01(*n A* is incorporated into the synaptic gating equations. The connectivity matrix *C* was multiplied by 0.2, and the Euler–Maruyama method (integration step *dt* = 1) was used for the numerical integration.

**Table 1.**
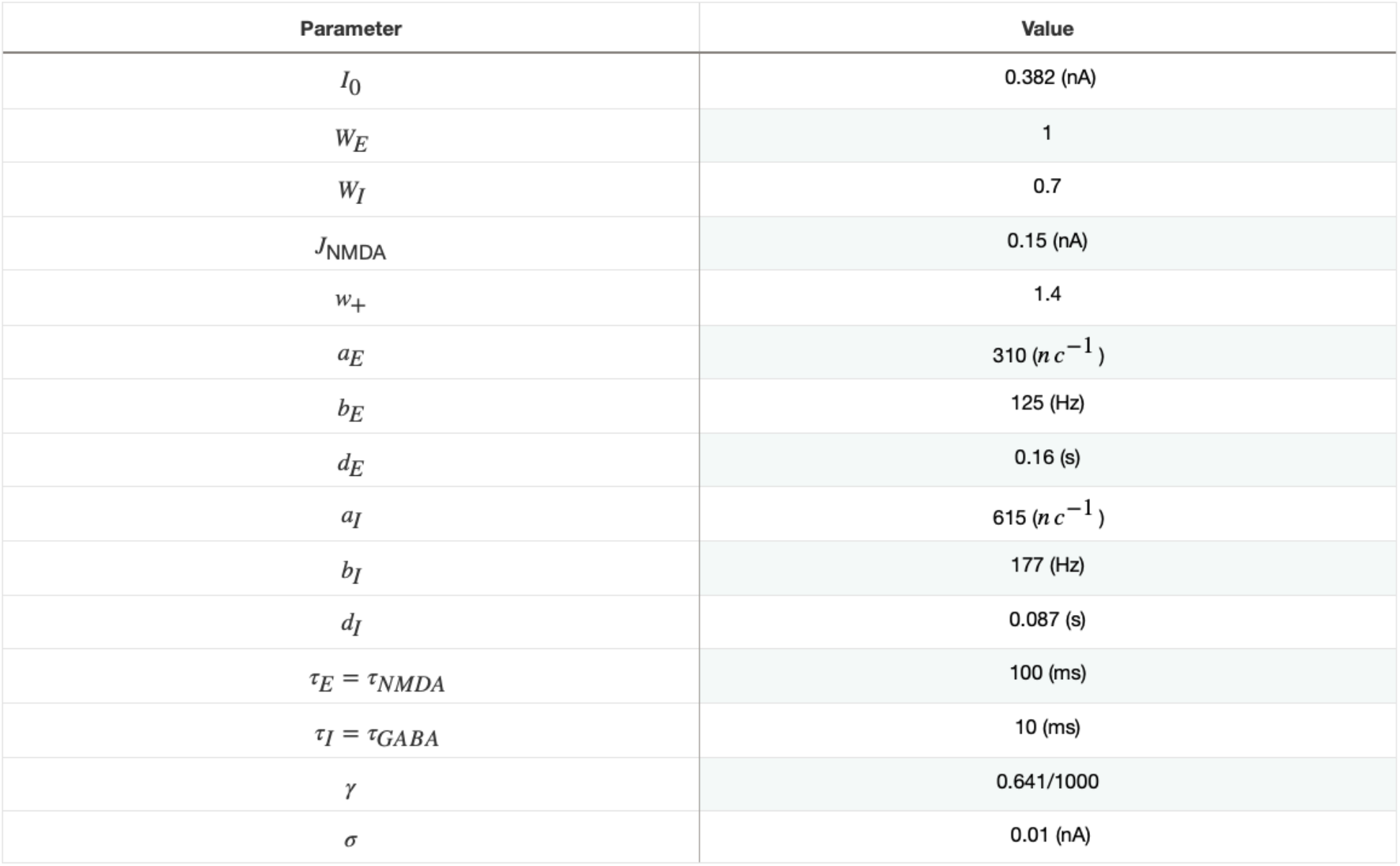
Biophysical parameters of the BEI model.

| Parameter | Value |
| --- | --- |
| $I_0$ | 0.382 (nA) |
| $W_E$ | 1 |
| $W_I$ | 0.7 |
| $J_{NMDA}$ | 0.15 (nA) |
| $w_+$ | 1.4 |
| $a_E$ | 310 ($nc^{-1}$) |
| $b_E$ | 125 (Hz) |
| $d_E$ | 0.16 (s) |
| $a_I$ | 615 ($nc^{-1}$) |
| $b_I$ | 177 (Hz) |
| $d_I$ | 0.087 (s) |
| $\tau_E = \tau_{NMDA}$ | 100 (ms) |
| $\tau_I = \tau_{GABA}$ | 10 (ms) |
| $\gamma$ | 0.641/1000 |
| $\sigma$ | 0.01 (nA) |

### Introducing receptor map heterogeneity

We optimized our patient-specific whole-brain simulation with the goal of determining the regional excitability values *A*^*exc*^ that best reproduced the empirical Functional Connectivity (FC) and Functional Connectivity Dynamics (FCD). To estimate *A*^*exc*^, we differentiated between cortical 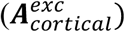 and subcortical regions 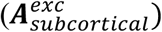, as receptor density maps were only available for cortical areas.

For cortical regions, excitability was modelled as a linear combination of receptor densities, receptor-specific scaling factors and a constant bias term. Specifically, we defined a cortical receptor matrix 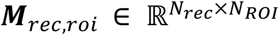, containing the normalized densities of *N*_*rec*_ receptor types across *N*_*ROI*_ cortical regions, and a vector of receptor scaling parameters 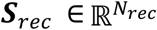. The cortical excitability was then computed as:

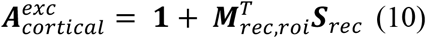

where 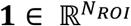 is a vector of ones representing the bias term. Importantly, ***S***_*rec*_ was independently estimated with a particle swarm optimization (PSO; see later *Model Optimization*) for perilesional and non-perilesional areas leading to 38 parameters (i.e., 19 receptor scalings for each 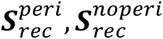). In subcortical regions, 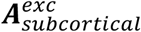 was directly optimized as two independent free parameters corresponding to perilesional and non-perilesional areas, unconstrained by receptor densities. This receptor-based formalism provides a dual insight: it enables the quantification of differential receptor contributions to regional excitability in perilesional and non-perilesional cortex and allows the reconstruction of individualized excitability profiles across the entire brain.

### Hemodynamic Model

We generated the BOLD signal from the simulated excitatory firing rate 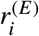 by transforming it with the hemodynamic Balloon–Windkessel model (Stephan et al., 2007). Firstly, the excitatory firing rate together with an auto-regulatory feedback determine vasodilator signal *s*_*i*_. This results in proportional changes in blood flow *f*_*i*_, which consequently affect blood volume *v*_*i*_ and deoxyhaemoglobin levels *q*_*i*_. Specifically, this process is described by:

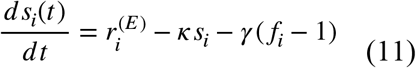

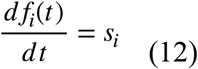

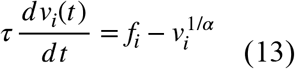

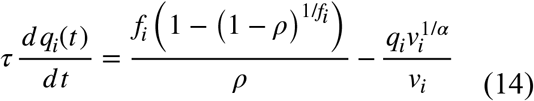

with *γ* representing the rate constant for autoregulatory feedback by blood flow, *κ* the rate constant of the vasodilatory signal decay, the resistance of the veins by *α, τ* as a time constant, and *ρ* as the resting oxygen extraction fraction. The BOLD signal for each brain region *B*_*i*_ is given by the blood volume *v*_*i*_ and the deoxyhaemoglobin content *q*_*i*_, described as:

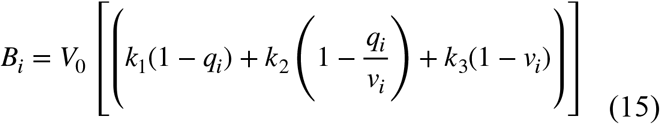

A detailed account of the hemodynamic model and all associated biophysical parameters is available in Stephan et al. (2007) and Supplementary Table ST7. Consistent with the pre-processing of the empirical data, the simulated BOLD time series were subjected to global signal regression, linear detrending, and band-pass filtering between 0.008 and 0.08 Hz.

### Functional Connectivity and Functional Connectivity Dynamics

For both empirical and simulated BOLD data, functional connectivity (FC) was quantified by computing the Pearson correlation between all pairs of brain regions. To capture the temporal evolution of these connectivity patterns, we computed the Functional Connectivity Dynamics (FCD). This metric was obtained using a sliding-window approach with a window length of 60 TR and a step size of 12 TR. Within each window, we calculated the instantaneous FC; subsequently, these matrices were correlated across all time windows to generate the final FCD matrix.

### Model Optimization

To fit the patient-specific brain models, we followed a two-step optimization procedure with the overall goal of best approximating the empirical FC and FCD. Model performance was quantified using an error metric defined as: 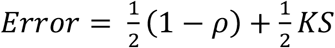, where *ρ* denotes the Pearson correlation coefficient between empirical and simulated FC, and *KS* reflects the Kolmogorov–Smirnov distance between the empirical and simulated FCD distributions.

In the first step, we optimized the global coupling parameter G by systematically sampling values from 2 to 12.5 in increments of 0.025 and identified the optimal G for each patient. In the second step, we applied a particle swarm optimization (PSO; https://pythonhosted.org/pyswarm/) to optimize the perilesional (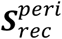; 19 parameters) and non-perilesional (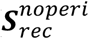; 19 parameters) receptor scaling parameters, together with the prelesional and non-perilesional subcortical excitability values (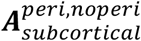; 2 parameters). We constrained the regional excitability values *A*^*exc*^ to remain within the range of 0.96 to 1.04 during optimization, and the NMDA receptor effects were restricted to non-negative (excitatory) while GABA-A receptor effects were restricted to non-positive (inhibitory) contributions. For each subject, we employed a multi-swarm optimization approach consisting of 10 swarms, each containing 60 particles and running for a maximum of 9 iterations. The best parameter estimate across all swarms was then selected for that subject. This PSO optimization framework resulted in a good model fit (see Fig. S1a-c).

### Behavioural outcome regressions

To examine how receptor scalings explain acute and long-term behavioural outcomes, we applied ridge regression (Hoerl & Kennard, 1970) within a leave-one-out cross-validation (LOOCV) paradigm. Ridge regression applies an L2 regularization penalty to stabilize coefficient estimates in the presence of multicollinearity and high-dimensional predictors. The model included lesion-related covariates such as lesion etiology (ischemic vs. hemorrhagic), lesion location, and regional structural connectivity (SC) damage. SC damage was represented as nodal white matter pathway damage and reduced in dimensionality using PCA, retaining five component scores to explain approximately 65% of the variance. The ridge regularization parameter (*α*) was optimized using a logarithmically spaced grid (10^−5^ – 10^5^), selecting the value minimizing cross-validated mean squared error (*MSE*). Model performance was quantified using LOOCV-based MSE and coefficient of determination (*R*^*2*^). Final coefficient estimates were obtained by averaging standardized coefficients across all LOOCV folds at the optimal *α*. Model and coefficient significances were assessed via permutation testing (10,000 permutations), with p-values corrected using the Benjamini–Hochberg false discovery rate (FDR) procedure.

To assess how hub-region excitability relates to behavioural outcomes, we fitted multivariate linear regression models with lesion etiology as a covariate. These models explicitly tested interactions between nodal excitability and its corresponding structural pathway damage to assess whether the behavioural relevance of excitability depended on the degree of structural disconnection. Non-significant interaction terms were removed. Structural damage outside hub regions was summarized using four principal component scores (∼65% variance explained). Standardized regression coefficients and FDR-corrected p-values were reported, with correction for multiple comparisons applied to the target variables of interest, excluding the intercept and lesion etiology covariate. Post hoc simple slope analyses were conducted for significant interactions.

## Results

### Fitting neurochemically informed digital twins

To assess how the neurochemical architecture after a stroke shapes behavioural outcome, we fitted patient-specific digital twins (i.e., whole-brain model) with their own neuromodulatory profiles to fMRI data from 96 acute stroke patients (Fig. 1). This allowed us to derive subject-wise regional cortical excitabilities along with perilesional and non-perilesional receptor scaling parameters for 19 receptor and transporter density maps. These scaling parameters quantify each map’s contribution to neuronal excitability and may reflect direct as well as indirect biological mechanisms, with indirect effects being particularly likely for transporters. NMDA scalings were constrained to be non-negative and GABA-A scalings to be non-positive, consistent with their excitatory and inhibitory roles. We optimized the global coupling parameter first (Model Error M = 0.365, SD = 0.028, N = 96), and afterwards the receptor scaling parameters together with the subcortical excitability values (Model Error M = 0.329, SD = 0.031), resulting in a significant model fit improvement (Wilcoxon signed-rank test, one-sided; W = 0.0, p < 0.001, N = 96; see supplementary S1c). To assess whether model performance depended on the spatial topology of the receptor maps rather than on spatial autocorrelation (SA), we repeated the model fitting using spatially autocorrelated surrogate maps, which yielded significantly poorer fits (Wilcoxon signed-rank test, one-sided; W = 1300, p < 0.001, N = 96; see supplementary S1d).

### Acute motor performance explained by multi-receptor profile

After fitting the digital twins to the acute-stage empirical data, we firstly examined the distributions of the estimated receptor scaling parameters (Fig. 2a). Interestingly, Histaminergic H3 scaling showed a marked spatial dissociation, with substantially lower values in perilesional compared to non-perilesional regions (median –0.002 vs 0.010; Fig. 2b). This within-patient difference was significant (Wilcoxon signed-rank test, W = 1065, N = 90, p < 0.001) and remained after false discovery rate correction (*p(FDR)* = 0.002). No other receptor scaling remained significant after correction. These results indicate that the effect of histaminergic H3 receptors on neuronal excitability differs between affected regions.

**Figure 2.**
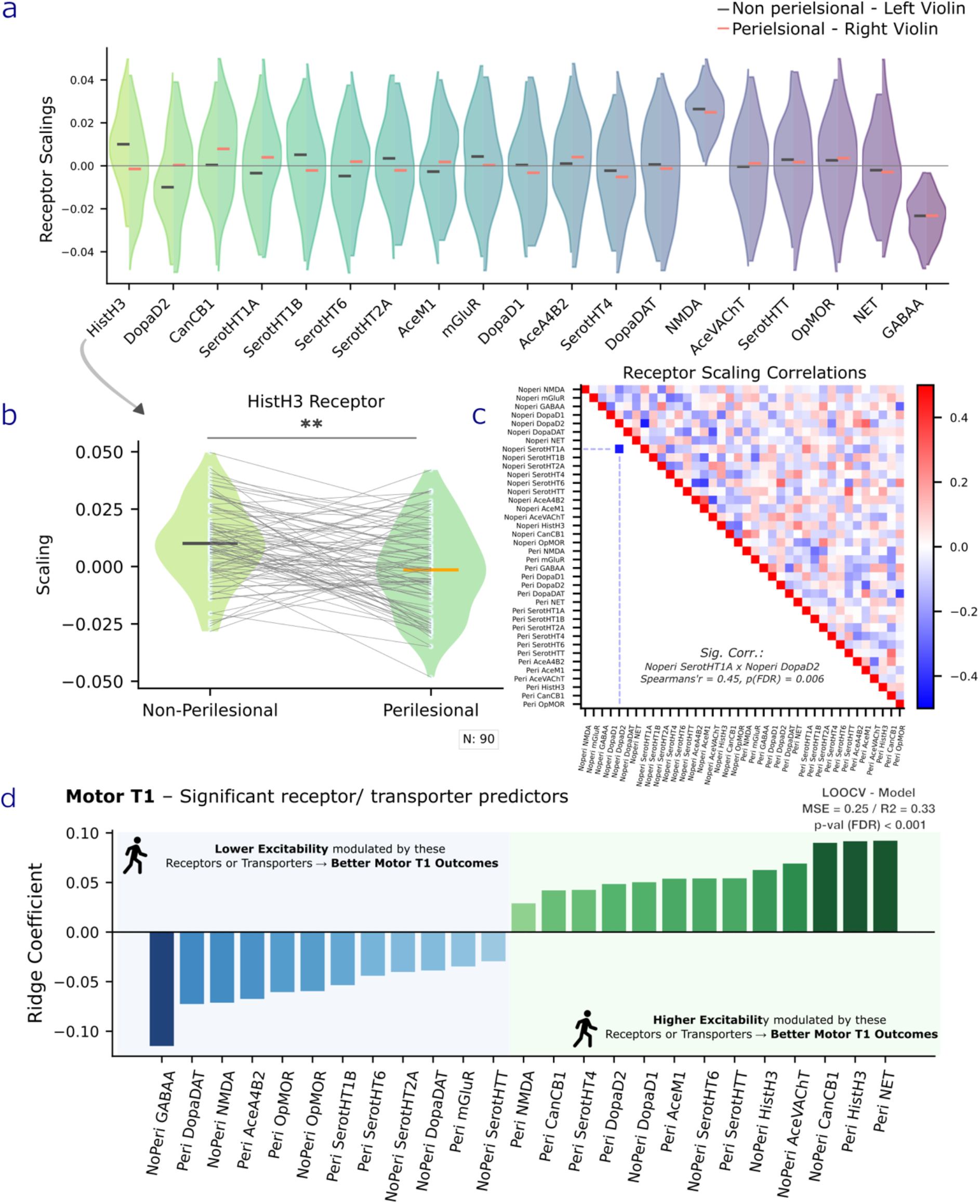
Receptor and transporter scaling characterization and predictors of acute (T1) motor outcomes. **a**| Distribution of model-derived receptor scaling parameters across perilesional and non-perilesional cortical regions. Each violin represents the fitted scaling of a given receptor or transporter, with left violins corresponding to non-perilesional and right to perilesional regions. Median values are indicated as horizontal lines. Positive or negative values correspond to increased or decreased neuronal excitability, respectively. **b**|Histamine H3 (HistH3) receptor scalings differed significantly between perilesional and non-perilesional regions, suggesting region-specific modulation of neuronal excitability. Grey lines connect within-patient values. **c**|Spearman correlation matrix of receptor scaling parameters across patients, separately for perilesional and non-perilesional regions. **d**|Significant coefficients of acute motor performance (T1) identified using a ridge regression leave-one-out cross-validation (LOOCV) framework with subsequent permutation testing. Non-perilesional receptor GABA-A, cannabinoid CB1, perilesional HistH3, and norepinephrine transporter (NET) scalings emerged as leading predictors of motor outcomes. The shaded green background highlights receptors or transporters that associate with better motor outcomes when they positively modulate excitability, while the blue area highlights variables that associate with improved outcomes when they negatively modulate excitability.

To further characterize the relationship between perilesional and non-perilesional receptor scalings, we examined their pairwise correlations (Fig. 2c). On average, the Spearman correlation was close to zero, *r*_*s*_ = −0.02, but exhibited variability (*SD* = 0.12). We found a significant anticorrelation between non-perilesional DopaD2 and non-perilesional serotonin HT1A receptor scalings (*r*_*s*_ *=* −0.45, *N* = 90, *p(FDR)* = 0.006; see also supplementary Fig. S2). To interpret this anticorrelation, we examined the underlying DopaD2 and serotonin HT1A receptor maps and found that they were highly correlated (*r*_*s*_ *=* 0.84, *N* = 200, *p* < 0.001). This suggests that the opposing receptor scalings may reflect a compensatory mechanism balancing excitation.

To understand how the patient-specific neuromodulatory profiles are predictive of post-stroke behavioral outcomes, we examined how they explain motor, language, and general cognitive performances at the acute (T1) and one-year (T3) follow-up stages. We used ridge regression within a leave-one-out cross-validation (LOOCV), estimating the statistical model and coefficient significance via permutation testing. We controlled for stroke localization, lesion type, and five principal components capturing regional structural connectivity damage (see Methods). After correction for multiple comparisons, all regression models except Cognition T1 were significant (Supplementary Fig. S3a; Supplementary Table ST1). Significant scaling coefficients were observed in the regression models explaining acute (T1) motor and language outcomes (Fig. 2d, 3a; Supplementary Table ST2,3).

The regression model for acute motor scores was highly significant (*N* = 78; *R*^*2*^ = 0.33; *MSE* = 0.25; FDR-corrected model *p* < 0.001) and identified multiple significant scaling coefficients (Fig. 2d). Positive coefficients (Fig. 2d; green box) indicate that positive receptor scalings—i.e., higher excitability modulated by the respective receptors or transporters—were associated with better motor outcomes, and conversely, that negative scalings were linked to worse outcomes. The strongest positive coefficients were observed for non-perilesional CB1 receptor, perilesional H3 receptor, and norepinephrine transporters, suggesting that a higher excitability modulated by these receptors and transporter was associated with better motor outcomes. Interestingly, both perilesional and non-perilesional cannabinoid CB1 and histaminergic H3 exhibited positive coefficients. In contrast, NMDA, serotonin transporter, and serotonin HT6 receptor coefficients showed location-dependent associations with motor outcomes, differing between perilesional and non-perilesional areas. This asymmetry highlights the importance of distinguishing between tissue nearby or distant to the stroke.

Negative ridge coefficients (Fig. 2d; blue box) indicate that negative scalings—i.e., lower excitability modulated by the respective receptors or transporters—were associated with better motor outcomes, and conversely, that positive scalings were linked to worse outcomes. Among all coefficients, non-perilesional GABA-A emerged as the strongest (negative) coefficient. As GABA-A receptors are inhibitory, this indicates that more GABA-A-driven inhibition was positively associated with acute motor outcomes, and conversely, disinhibition was linked to poorer outcomes. Likewise, the coefficient for non-perilesional NMDA was negative; since NMDA is an excitatory receptor, this demonstrates that better outcomes were driven specifically by a decrease in non-perilesional NMDA-modulated excitation. Among other significant predictors, reduced excitability modulated by dopamine transporters or perilesional α4β2 nicotinic acetylcholine receptors was also markedly associated with improved motor outcomes. Together, these results indicate that acute motor outcomes are driven by how various receptors and transporters modulate excitability across perilesional and non-perilesional regions.

## Common receptor clusters linked to acute language and motor performance

The regression model explaining acute language performance (Fig. 3a; *N* = 84; *R*^*2*^ = 0.25; *MSE* = 0.58; FDR-corrected model *p* < 0.001) also revealed significant receptor scaling coefficients. NMDA- and acetylcholine transporter–modulated excitability showed strong and asymmetric effects across perilesional and non-perilesional regions on language performance (Fig. 3a; Supplementary Table ST3). Also, lower excitability modulated by perilesional serotonin HT1B receptors was markedly associated with improved language outcomes.

**Figure 3.**
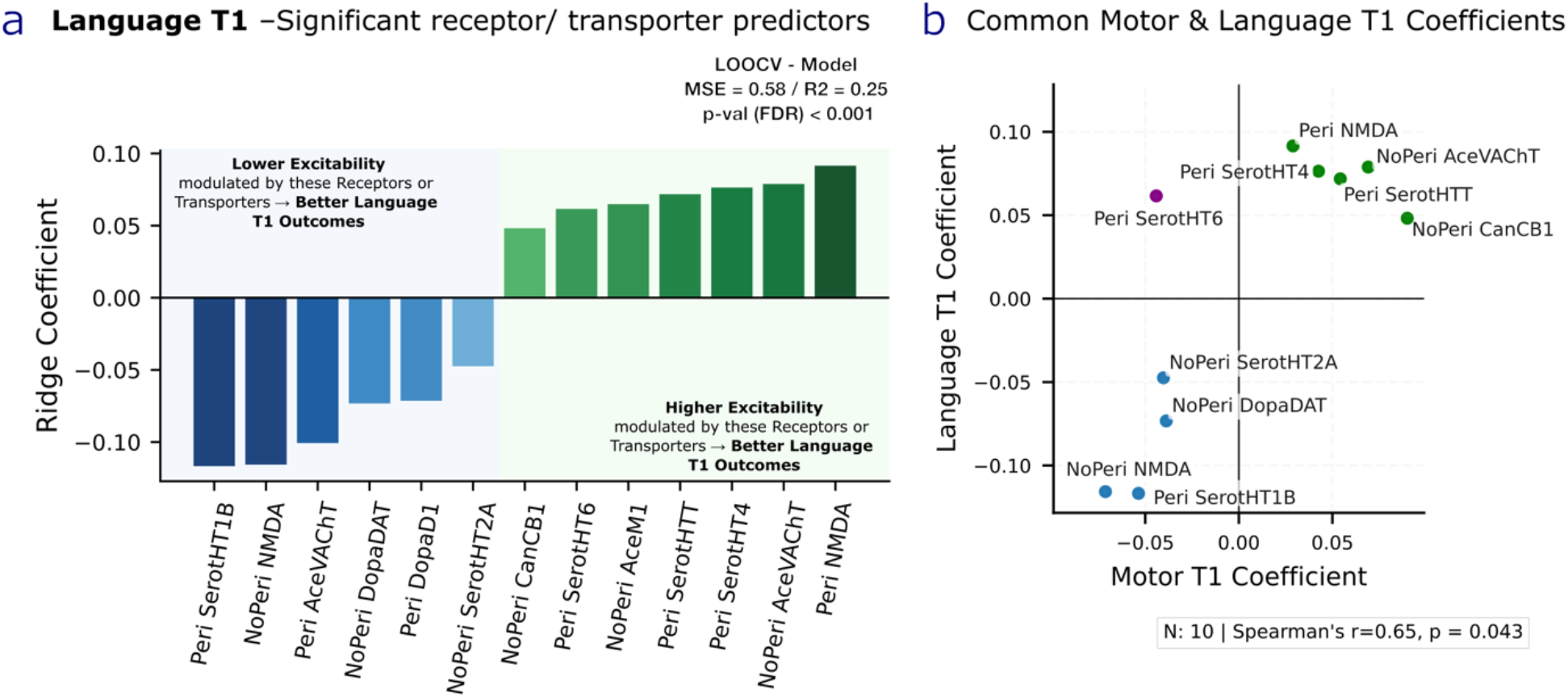
Significant receptor and transporter scaling coefficients for acute (T1) language outcomes and common behavioural receptor clusters. **a**|Significant scaling coefficients for acute language performance. Asymmetric effects of NMDA- and acetylcholine transporter–driven excitability changes in perilesional and non-perilesional regions on language performance can be observed. The shaded green background highlights receptors or transporters that associate with better motor outcomes when they positively modulate excitability, while the blue area highlights variables that associate with improved outcomes when they negatively modulate excitability. **b**|Common significant receptor and transporter coefficients for acute language and motor performance. Receptor and transporter scaling coefficients showed broadly similar effects across both behavioural domains. Two distinct clusters emerged: one in which increased excitability was associated with better outcomes (upper right; green dots), and another in which decreased excitability corresponded to better behavioural performance (lower left; blue dots).

Most receptor and transporter scaling coefficients that were significant for acute language performance also determined acute motor outcomes, and their effect directions were broadly aligned (Fig. 3b; *Spearman’s r* = 0.65; *N* = 10; *p* = 0.043). Interestingly, two clusters emerged: in one group, receptor-map–modulated increases in perilesional and non-perilesional excitability were positively associated with acute behavioural outcomes (Fig. 3b, green dots), in the other, excitability decreases were linked to better performances (blue dots), potentially reflecting shared functional properties within each cluster. In contrast to these two clusters, an increase in excitability by perilesional serotonin HT6 receptors was positively associated with language and negatively with acute motor performance.

### Cortical hub excitabilities determine acute and long-term behavioural outcomes

Our brain modelling framework allowed us to quantify the excitability across all cortical brain regions in individual stroke patients (ROIs; Fig. 4a). To explore the relationship between structure and regional excitability, we correlated each cortical region’s mean excitability across patients with its structural connectedness (i.e., degree of the average structural connectivity matrix; see Methods). We observed a significant anticorrelation (Fig. 4b; *Spearman’s r* = –0.38; *N* = 200; *p* < 0.001), indicating that generally highly connected regions tended to exhibit lower excitability, and vice versa. We next examined how regional connectivity damage relates to excitability in individual patients by analysing structurally damaged regions and their corresponding excitability values. To account for excitability differences driven by a region’s degree, we normalized each region’s excitability by its node degree and quantified damage as the cumulative loss of healthy connection strength per region. This analysis revealed a significant anticorrelation (*Spearman’s r* = −0.23; *N* = 7339; *p* < 0.001; Fig. 4c), indicating that regions with greater structural connectivity damage exhibited reduced excitability.

**Figure 4.**
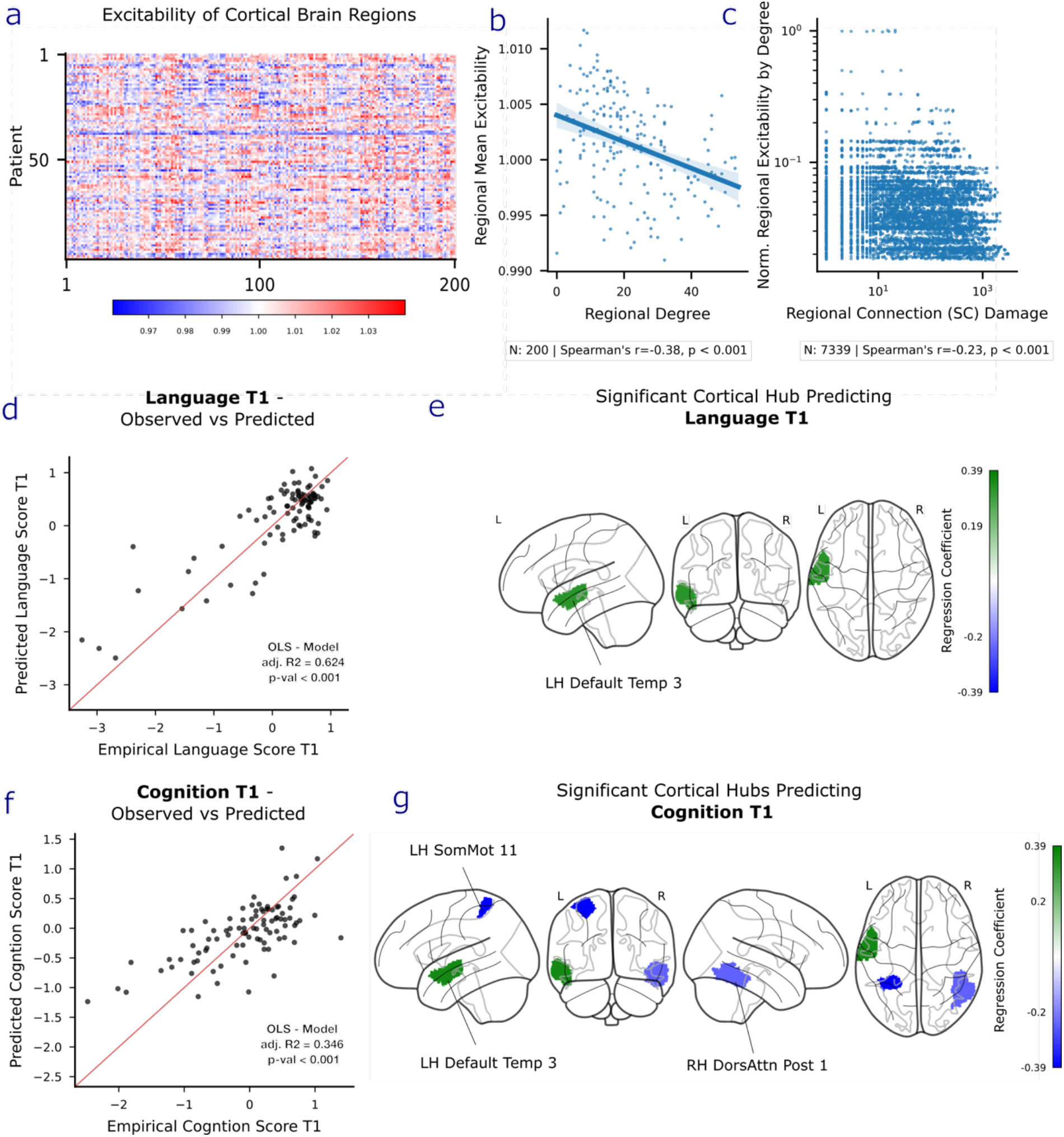
Cortical hub excitabilities determine acute cognitive as well as acute and long-term language scores. **a**|Heatmap illustrating the patient-specific excitability profiles for all cortical brain regions. Warmer colors indicate increased excitability. **b**|Relationship between mean regional excitability over all patients and regional structural connectedness (i.e., degree). Brain regions with a higher degree generally exhibited a lower excitability and vice versa. **c**|We next assessed the relationship between regional structural connectivity (SC) damage and excitability. Normalizing excitability by node degree, as motivated by the earlier finding, revealed that regions with higher SC damage exhibited lower excitability. **d** | Regression model performance for acute language outcomes, illustrating the relationship between empirical and predicted language T1 scores. **e**|Brain visualization of significant cortical hub excitability predicting acute language performance. LH Default Temp 3 emerged as a significant positive predictor. **f** | Model performance for acute cognition outcomes. **g** | Brain visualization of significant cortical hub excitabilities predicting acute cognitive performance (T1). LH Default Temp 3 was again a positive predictor, and LH SomMot 11, as well as RH DorsAttn Post 1, negatively predicted acute scores.

Previous research has shown that the structural integrity of brain hubs plays an important role in long-term recovery (Aben et al., 2019; Ktena et al., 2019; Schirmer et al., 2019; Stam, 2024). We therefore investigated whether hub excitability, particularly as modulated by damage, also influences post-stroke behavioural outcomes. We focused on the five most connected hubs, defined by nodal degree, and examined their association with behavioural scores using linear regression models. The models accounted for lesion etiology (ischemic versus hemorrhagic) and overall non-hub nodal damage, captured via principal components of nodal damage. For each hub, we explicitly tested interactions between regional excitability and local nodal hub damage (see Methods).

Acute language performance (*N* = 90) was robustly explained by the regression model, accounting for a substantial proportion of variance (adjusted *R*^*2*^ = 0.62; *F*(15,74) = 5.30, *p* < 0.001; Fig. 4d). Interestingly, higher excitability in the left default temporal region 3 was positively associated with acute language performance (*β* = 0.32, *z* = 2.95, *p* = 0.003, *p(FDR)* = 0.045; Fig. 4e) and remained the only significant predictor after FDR correction (see supplementary table ST4 for all predictors). In contrast, excitability of the right dorsal attention network (posterior) node 1 was negatively associated with acute language scores (*β* = −0.33, *z* = −2.39, *p* = 0.017, *p(FDR)* = 0.118), although this effect did not remain significant after correction for multiple comparisons. This result indicates that preserved acute language performance is associated with increased excitability in the left temporal default region 3.

The regression model explaining acute cognitive performance (N = 92) was also significant (adjusted *R*^*2*^ = 0.35; *F*(16,75) = 4.01, *p* < 0.001; Fig. 4f) and revealed several highly significant hub-specific predictors (see supplementary table ST5 for all predictors). As observed in the language domain, excitability in the left default temporal region 3 was again positively associated with performance (*β* = 0.36, *t* = 3.68, *p* < 0.001, *p(FDR)* = 0.002; Fig. 4g). Also, in the right hemisphere, increased excitability in the posterior dorsal attention node 1 was negatively associated with cognition (*β* = -0.25, *t* = -2.39, *p* = 0.020, *p(FDR)* = 0.049). We found a significant interaction between excitability and nodal damage in the left somatomotor region 11 (*β* = 0.2, *t* = 3.12, *p* = 0.003, *p(FDR)* = 0.008). A simple slope analysis revealed that increased excitability strongly predicted lower cognitive performance in patients with no nodal damage (*β* = −0.48, *t* = −4.14, *p* < 0.001) and average damage (*β* = −0.39, *t* = −3.86, *p* <0.001). This negative relationship was substantially attenuated in patients with high structural damage (*β* = −0.19, *t* = −2.03, *p* = 0.05; Supplementary Fig. S4). These results demonstrate that acute cognitive performance mirrors language function, specifically the favourable role of the left temporal default region 3 and the negative association of the right dorsal attention node excitability. However, acute cognition also implicates the left somatomotor network ROI (11), where higher excitability was associated with impairment, especially in patients with non-severe structural hub damage.

Lastly, with respect to one year post-stroke language recovery, we found that the regression model (*N* = 58) explained a substantial proportion of variance (adjusted *R*^*2*^ = 0.68; *F*(16,41) = 8.72, *p* < 0.001; Fig. 5a). Beyond the strong predictive value of initial severity (Language T1: *β* = 0.31, t = 6.45, *p* < 0.001, *p(FDR)* < 0.001; supplementary table ST6 for all predictors), we also identified the acute excitability of the right posterior dorsal attention node 1 as a crucial predictor of long-term language recovery. Although increased excitability in this region has been associated with poorer outcomes in the acute stage, it demonstrated here a significant compensatory role for long-term recovery (*β* = 0.16, *t* = 2.82, *p* = 0.007, *p(FDR)* = 0.037; Fig. 5b). This effect was qualified by a significant interaction with nodal damage (*β* = 0.20, *t* = 3.24, *p* = 0.002, *p(FDR)* = 0.018). A simple slopes analysis (Fig. 5C) revealed that higher acute nodal excitability was significantly associated with better language outcomes in patients with high nodal damage (+1SD; *β* =0.36, *t* =3.66, *p* < 0.001) and average nodal damage (Mean; *β* = 0.16, *t* = 2.82, *p* = 0.01), but not in patients with no nodal damage (*Raw score* = 0, *Z* = −0.44; *β* =0.08, *t* = 1.41, *p* = 0.17). These results show that long-term language recovery is predicted by higher acute excitability of the right dorsal attention hub, particularly in patients with greater structural hub damage.

**Figure 5.**
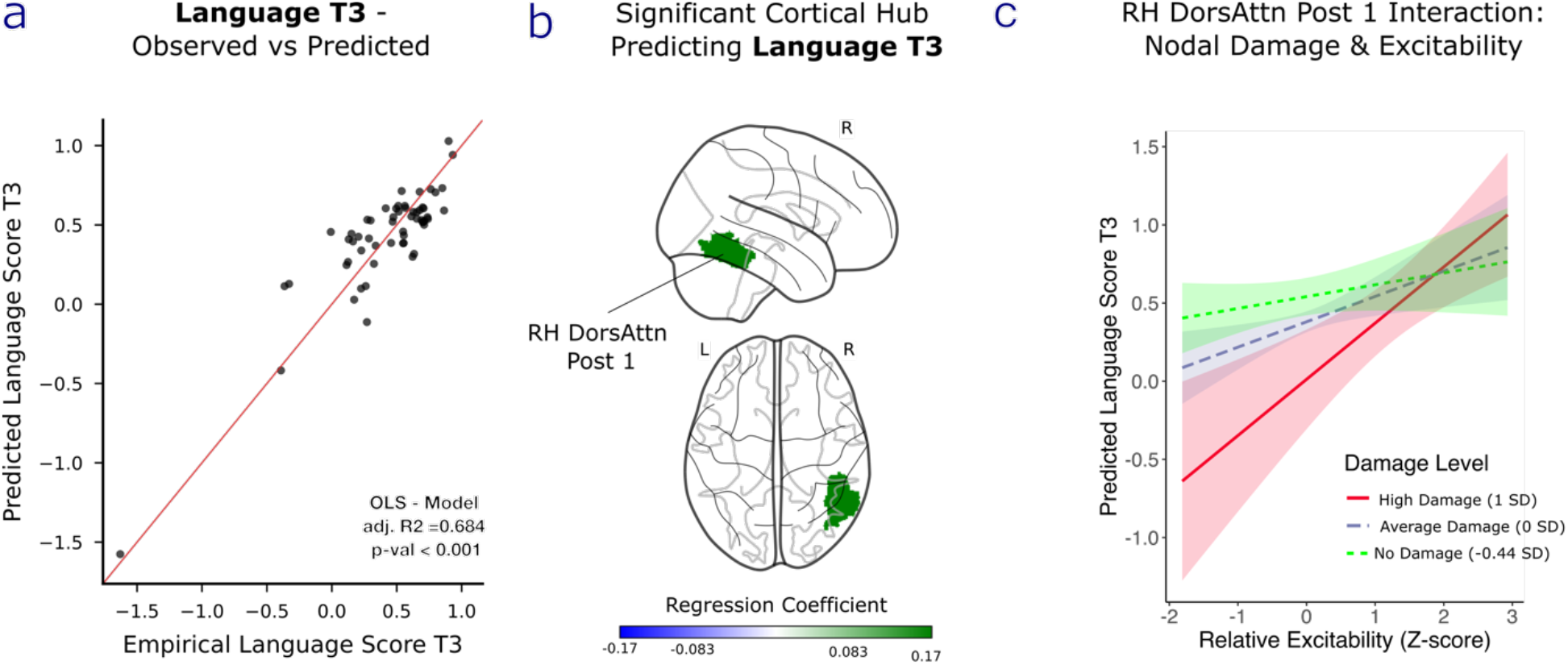
Long-term language recovery predicted by the interaction of acute cortical hub excitability with hub damage. **a** | Scatter plot showing significant prediction of T3 language scores by the regression model. **b** | Brain visualization of significant acute cortical hub excitability predicting long-term language performance (T3). RH DorsAttn Post 1 was a positive predictor for long-term scores. **c** | Simple slope plots show that higher acute hub excitability significantly predicts better language outcomes in patients with high (+1 SD, red) and average (mean, blue) damage. In contrast, this association is absent in patients with no nodal damage (-0.44 SD, green). The shaded areas indicate the confidence intervals.

## Discussion

The principal aim of this study was to employ patient-specific digital twin to estimate how receptor and transporter systems modulate cortical excitability and, in turn, explain patient outcomes. We found that acute motor performance was explained by the excitability modulation of various receptors and transporters: higher inhibition by non-perilesional GABA-A receptors and increased excitability modulated by non-perilesional CB1 receptors, perilesional histaminergic H3 receptors, and norepinephrine transporters were most strongly linked to favourable outcomes. Similar patterns were observed for acute language performance, and shared predictors across both domains formed two clusters in which increased excitability was either beneficial or detrimental. At the regional level, highly connected and structurally damaged regions exhibited reduced excitability. Interestingly, the excitability of structural brain hubs emerged as a key predictor of acute language and cognitive outcomes, and importantly, for long-term language recovery. Our findings demonstrate that receptor- and transporter-driven modulation of cortical excitability, together with the excitability of cortical hubs, determine post-stroke behavioural outcomes.

We found that various estimated receptor scalings, which capture the impact of receptors and transporters on regional excitabilities, were significant predictors of acute motor scores. Non-perilesional GABA-A excitability most strongly explained acute motor performance, with higher inhibition associated with better outcomes and greater disinhibition with worse outcomes. The important role of GABA-A receptors in stroke patients was previously underlined by Kim et al. (2014), who reported that a higher GABA-A receptor availability, as a measure of neuronal density and integrity, at approximately one month post-stroke, positively correlates with both contemporaneous and 3-month motor performance. Also, we found that increased NMDA-driven excitability in non-perilesional regions was associated with reduced acute motor outcomes. These findings suggest that unspecific, i.e., non-perilesional increases in excitability within the core excitatory–inhibitory system are linked to worse motor performance in the acute phase.

In contrast, higher perilesional NMDA scalings were associated with improved motor outcomes, highlighting the differential role the same neuroreceptor can have depending on the stroke lesion proximity. These findings add to the complex role of NMDARs after stroke. While it is well established that early NMDAR overactivation triggers excitotoxicity, accumulating evidence suggests that restoring or normalizing NMDAR signalling at later stages can support neuronal survival and recovery (Ge et al., 2020). It remains to be determined whether such neuroprotective effects primarily depend on timing, engagement of specific NMDAR subtypes, or receptor localization (Ge et al., 2020). Next to these factors, our results suggest that the functional impact of NMDAR-related excitability also critically depends on lesion proximity.

Elevated excitability modulated by perilesional norepinephrine transporter and histaminergic H3 receptors, together with non-perilesional cannabinoid CB1 receptors, emerged as the strongest positive drivers of acute motor outcomes. Notably, the modulation of receptor and transporter maps to cortical excitability may reflect both direct and indirect underlying biological mechanisms, the latter being particularly plausible for transporters. Also, perilesional cannabinoid and non-perilesional histaminergic receptor scalings were likewise positive predictors, indicating a consistent unidirectional effect across these receptor maps. For cannabinoid CB1 receptors (CB1R), previous research has shown that agonists can shift the excitation-to-inhibition balance towards excitation (Den Boon et al., 2015), and neuroprotective effects have been observed after pharmacological intervention. Counterintuitively, both CB1R agonists and antagonists have been reported to improve stroke outcomes (Bietar et al., 2023; Carter et al., 2024). Proposed mechanisms include attenuation of excitotoxicity, enhanced neurogenesis, upregulation of neuroprotective factors, modulation of inflammatory responses, and the activation of specific cell signalling pathways (Bietar et al., 2023). Beyond these findings, our results indicate that CB1R-driven excitability changes in both affected and non-affected regions may be critical for acute motor outcomes, highlighting the potential pharmacological relevance of CB1R-mediated excitability.

Regarding histaminergic receptors, animal models of ischemic stroke have shown that after neuronal injury, a rising level of histamine can be observed together with increased H3 receptor binding densities in cortical and subcortical brain regions (Hu & Chen, 2012; Lozada et al., 2005). We found that histaminergic H3 receptors in non-perilesional brain regions exerted a greater influence on neuronal excitability than those in perilesional regions; however, increased scalings in both regions were positively associated with improved motor outcomes. This suggests that generally enhanced histaminergic receptor–driven excitability is linked to better acute motor outcomes. Nevertheless, the mechanisms by which H3 receptors modulate neuronal excitability are complex and likely depend on multiple localized factors (Yang et al., 2022). Beyond these receptors, multiple additional receptor and transporter scalings emerged as significant predictors, underscoring the complex landscape through which these maps shape cortical excitability and, in turn, acute motor outcomes.

Receptor and transporter scalings were also explanatory for acute language outcomes. We found contrastive roles of receptor and transporter scalings depending on their lesion proximity. Perilesional NMDA scalings emerged as the leading predictor of acute language outcomes, whereas non-perilesional NMDA scalings were negatively associated with acute language outcomes. Likewise, cholinergic transporters showed a positive association with acute language outcomes in non-perilesional regions but a negative association in perilesional regions. This finding is notable as acetylcholine is involved not only in attention and memory but also because its diffuse cortical projections target language-related areas (Katsari et al., 2022). These cholinergic innervations have been proposed to underlie potential beneficial effects reported for cholinergic pharmacological interventions in post-stroke aphasia (Katsari et al., 2022). Nevertheless, our findings hint at the heterogenous roles of perilesional and non-perilesional NMDAR as well as cholinergic transporters, which could be insightful for future drug development.

Other receptor scalings were also significant for the language domain, exhibiting a multifaceted impact of these neuromodulatory systems. Despite this apparent complexity, the analysis of predictors common to both domains interestingly revealed two distinct clusters: In one cluster, receptor map–induced increases in perilesional and non-perilesional excitability were associated with improved acute behavioural outcomes, whereas in the other cluster, comparable increases in excitability were associated with poorer performance. This finding indicates potential shared roles of these receptor groups and might be interesting targets for pharmacological investigations.

Our whole-brain simulation framework also allowed us to estimate the excitabilities of cortical brain regions in individual patients. We found that highly connected regions exhibit lower excitability than regions with fewer structural connections. This finding adds to previous research showing that the dynamics of structural hubs are more stable in healthy subjects (López-González et al., 2021) and that generally highly connected nodes exhibit reduced activity variance and susceptibility to perturbations (Ponce-Alvarez, 2025). Structural hubs are recognized as integrative cores of large-scale brain networks (Gong et al., 2009; Hagmann et al., 2008; Kaiser et al., 2007; Schulte et al., 2025; Sporns et al., 2007; Zamora-López et al., 2009, 2010, 2011) and crucial substrates of cognitive function in both health and disease (Crossley et al., 2014; Stam, 2024; Van Den Heuvel & Sporns, 2013). While lesion involvement and structural plastic changes of hub regions are important predictors of long-term behavioural recovery (Aben et al., 2019; Chen et al., 2021; Ktena et al., 2019; Schirmer et al., 2019; Stam, 2024; L. Wang et al., 2021), the influence of functional hub excitability has remained largely unexplored.

We found, that regarding acute outcomes, higher excitability in the left default temporal region 3 was associated with better language performance. Interestingly, while lateral temporal areas have been associated with key regions of the language network (Fedorenko et al., 2024; Lipkin et al., 2022), our findings were not specific to the language domain; excitability in this hub also served as a positive predictor of acute cognitive scores. This implies that the excitability of this hub plays a broadly supportive role for acute function. In contrast, higher excitability in the left somatomotor region 11 and the posterior right dorsal-attention region 1 was negatively associated with acute cognitive scores. Interestingly, the right dorsal-attention region 1 exhibited a distinct temporal dissociation. While associated with acute impairment, higher acute excitability in this region also strongly predicted favourable language outcomes at one-year post-stroke, specifically in patients with moderate-to-severe damage, but not in those with no local damage. This suggests a compensatory role: recruitment of this hub may be initially disruptive but becomes beneficial for long-term recovery when structural damage is moderate or worse. Therefore, this region represents a promising target for non-invasive brain stimulation, although a deeper understanding of its detailed compensatory role is required.

Despite these findings, the present study has several limitations. Although receptor scalings and regional hub excitabilities were explanatory, particularly for acute outcomes, associations with long-term recovery were more specific. While we identified a prominent role of a single hub for language recovery, we did not observe predictive hub excitabilities or receptor scalings for long-term motor or cognitive recovery. Also, cognitive outcomes in the current study were limited to measures of attention and memory. Future work should extend this approach to other cognitive domains. Also, we estimated receptor scalings separately for perilesional and non-perilesional areas. A further refinement could incorporate more detailed neurotransmitter disruption patterns, such as differential effects on pre- and postsynaptic neurons (Alves et al., 2025) and their consequences for neuronal excitability. With respect to the dataset, the generalizability of our model is constrained by the predominance of ischemic stroke in the cohort. Moreover, receptor scalings were estimated within the acute phase. Although this interval is relatively short, receptor expression may fluctuate during early post-stroke reorganization, potentially introducing additional variability. Finally, to reduce model complexity, we focused on the excitability of the excitatory neuronal population. While this marker impacts the excitatory-inhibitory ratio, future studies should also investigate the excitability of the inhibitory neuronal populations.

Taken together with these limitations, our results support the view that biophysically informed, patient-specific whole-brain modelling can mechanistically capture the influence of receptors and transporters on cortical excitability to explain acute deficits, while identifying integrative hub excitabilities as central drivers of functional outcomes across recovery timescales.

## Supporting information

Supplementary Materials Whole-brain receptor markers and hub excitabilities predict acute and long-term outcomes in stroke

## Data availability

Data can be made available upon request.

## Acknowledgments

The open source software FieldTrip (Oostenveld et al., 2011) was used for the visualization of cortical receptor density maps.

## References

Aben, H. P., Biessels, G. J., Weaver, N. A., Spikman, J. M., Visser-Meily, J. M. A., De Kort,P. L. M., Reijmer, Y. D., on behalf of the PROCRAS Study Group*, & Jansen, B. P.W. (2019). Extent to Which Network Hubs Are Affected by Ischemic Stroke Predicts Cognitive Recovery. Stroke, 50(10), 2768–2774. 10.1161/STROKEAHA.119.025637

Alia, C., Spalletti, C., Lai, S., Panarese, A., Micera, S., & Caleo, M. (2016). Reducing GABAA-mediated inhibition improves forelimb motor function after focal cortical stroke in mice. Scientific Reports, 6(1), 37823. 10.1038/srep37823

Alves, P., Nozais, V., Hansen, J. Y., Corbetta, M., Nachev, P., Martins, I. P., & Thiebaut De Schotten, M. (2025). Neurotransmitters’ white matter mapping unveils the neurochemical fingerprints of stroke. Nature Communications, 16(1), 2555. 10.1038/s41467-025-57680-2

Bietar, B., Tanner, S., & Lehmann, C. (2023). Neuroprotection and Beyond: The Central Role of CB1 and CB2 Receptors in Stroke Recovery. International Journal of Molecular Sciences, 24(23), 16728. 10.3390/ijms242316728

Caltana, L., Saez, T. M., Aronne, M. P., & Brusco, A. (2015). Cannabinoid receptor type 1 agonist ACEA improves motor recovery and protects neurons in ischemic stroke in mice. Journal of Neurochemistry, 135(3), 616–629. 10.1111/jnc.13288

Carter, C., Laviolette, L., Bietar, B., Zhou, J., & Lehmann, C. (2024). Cannabis, Cannabinoids, and Stroke: Increased Risk or Potential for Protection—A Narrative Review. Current Issues in Molecular Biology, 46(4), 3122–3133. 10.3390/cimb46040196

Chen, Y., Jiang, Y., Kong, X., Zhao, C., Zhong, S., Yang, L., Feng, T., Peng, S., Bi, Y., Corbetta, M., & Gong, G. (2021). Common and unique structural plasticity after left and right hemisphere stroke. Journal of Cerebral Blood Flow & Metabolism, 41(12), 3350–3364. 10.1177/0271678X211036606

Corbetta, M., Ramsey, L., Callejas, A., Baldassarre, A., Hacker, C. D., Siegel, J. S., Astafiev,S. V., Rengachary, J., Zinn, K., Lang, C. E., Connor, L. T., Fucetola, R., Strube, M., Carter, A. R., & Shulman, G. L. (2015). Common behavioral clusters and subcortical anatomy in stroke. Neuron, 85(5), 927–941. 10.1016/j.neuron.2015.02.027

Crossley, N. A., Mechelli, A., Scott, J., Carletti, F., Fox, P. T., McGuire, P., & Bullmore, E.T. (2014). The hubs of the human connectome are generally implicated in the anatomy of brain disorders. Brain, 137(8), 2382–2395. 10.1093/brain/awu132

Deco, G., Cruzat, J., Cabral, J., Tagliazucchi, E., Laufs, H., Logothetis, N. K., & Kringelbach, M. L. (2019). Awakening: Predicting external stimulation to force transitions between different brain states. Proceedings of the National Academy of Sciences, 116(36), 18088–18097. 10.1073/pnas.1905534116

Deco, G., Kringelbach, M. L., Arnatkeviciute, A., Oldham, S., Sabaroedin, K., Rogasch, N. C., Aquino, K. M., & Fornito, A. (2021). Dynamical consequences of regional heterogeneity in the brain’s transcriptional landscape. Science Advances, 7(29), eabf4752. 10.1126/sciadv.abf4752

Deco, G., Kringelbach, M. L., Jirsa, V. K., & Ritter, P. (2017). The dynamics of resting fluctuations in the brain: Metastability and its dynamical cortical core. Scientific Reports, 7(1), 3095. 10.1038/s41598-017-03073-5

Deco, G., Lynn, C. W., Sanz Perl, Y., & Kringelbach, M. L. (2023). Violations of the fluctuation-dissipation theorem reveal distinct nonequilibrium dynamics of brain states. Physical Review E, 108(6), 064410. 10.1103/PhysRevE.108.064410

Deco, G., Ponce-Alvarez, A., Hagmann, P., Romani, G. L., Mantini, D., & Corbetta, M. (2014). How local excitation-inhibition ratio impacts the whole brain dynamics. The Journal of Neuroscience: The Official Journal of the Society for Neuroscience, 34(23), 7886–7898. 10.1523/JNEUROSCI.5068-13.2014

Deco, G., Sanz Perl, Y., Vohryzek, J., Luppi, A. I., & Kringelbach, M. L. (2026).Neurotransmission-modulated whole-brain computation captures full task repertoire.Cell Reports, 45(1), 116816. 10.1016/j.celrep.2025.116816

Demirtaş, M., Burt, J. B., Helmer, M., Ji, J. L., Adkinson, B. D., Glasser, M. F., Van Essen, D. C., Sotiropoulos, S. N., Anticevic, A., & Murray, J. D. (2019). Hierarchical Heterogeneity across Human Cortex Shapes Large-Scale Neural Dynamics. Neuron, 101(6), 1181–1194.e13. 10.1016/j.neuron.2019.01.017

Dhawan, J., Benveniste, H., Luo, Z., Nawrocky, M., Smith, S. D., & Biegon, A. (2011). A New Look At Glutamate and Ischemia: Nmda Agonist Improves Long-Term Functional Outcome In A Rat Model of Stroke. Future Neurology, 6(6), 823–834. 10.2217/fnl.11.55

Fedorenko, E., Ivanova, A. A., & Regev, T. I. (2024). The language network as a natural kind within the broader landscape of the human brain. Nature Reviews Neuroscience, 25(5), 289–312. 10.1038/s41583-024-00802-4

Ge, Y., Chen, W., Axerio-Cilies, P., & Wang, Y. T. (2020). NMDARs in Cell Survival and Death: Implications in Stroke Pathogenesis and Treatment. Trends in Molecular Medicine, 26(6), 533–551. 10.1016/j.molmed.2020.03.001

Gong, G., He, Y., Concha, L., Lebel, C., Gross, D. W., Evans, A. C., & Beaulieu, C. (2009). Mapping Anatomical Connectivity Patterns of Human Cerebral Cortex Using In Vivo Diffusion Tensor Imaging Tractography. Cerebral Cortex, 19(3), 524–536. 10.1093/cercor/bhn102

Griffis, J. C., Metcalf, N. V., Corbetta, M., & Shulman, G. L. (2019). Structural Disconnections Explain Brain Network Dysfunction after Stroke. Cell Reports, 28(10), 2527–2540.e9. 10.1016/j.celrep.2019.07.100

Griffis, J. C., Metcalf, N. V., Corbetta, M., & Shulman, G. L. (2021). Lesion Quantification Toolkit: A MATLAB software tool for estimating grey matter damage and white matter disconnections in patients with focal brain lesions. NeuroImage. Clinical, 30, 102639. 10.1016/j.nicl.2021.102639

Hagmann, P., Cammoun, L., Gigandet, X., Meuli, R., Honey, C. J., Wedeen, V. J., & Sporns,O. (2008). Mapping the Structural Core of Human Cerebral Cortex. PLoS Biology, 6(7), e159. 10.1371/journal.pbio.0060159

Hansen, J. Y., Shafiei, G., Markello, R. D., Smart, K., Cox, S. M. L., Nørgaard, M., Beliveau, V., Wu, Y., Gallezot, J.-D., Aumont, É., Servaes, S., Scala, S. G., DuBois, J. M., Wainstein, G., Bezgin, G., Funck, T., Schmitz, T. W., Spreng, R. N., Galovic, M., … Misic, B. (2022). Mapping neurotransmitter systems to the structural and functional organization of the human neocortex. Nature Neuroscience, 25(11), 1569–1581. 10.1038/s41593-022-01186-3

Herzog, R., Mediano, P. A. M., Rosas, F. E., Luppi, A. I., Sanz Perl, Y., Tagliazucchi, E., Kringelbach, M. L., Cofré, R., & Deco, G. (2024). Neural mass modelling for the masses: Democratising access to whole-brain biophysical modelling with FastDMF. Network Neuroscience, 1–38. 10.1162/netn_a_00410

Hoerl, A. E., & Kennard, R. W. (1970). Ridge Regression: Biased Estimation for Nonorthogonal Problems. 12(1), 55–67.

Hu, W.-W., & Chen, Z. (2012). Role of Histamine and Its Receptors in Cerebral Ischemia. ACS Chemical Neuroscience, 3(4), 238–247. 10.1021/cn200126p

Idesis, S., Allegra, M., Vohryzek, J., Perl, Y. S., Metcalf, N. V., Griffis, J. C., Corbetta, M.,Shulman, G. L., & Deco, G. (2024). Generative whole-brain dynamics models from healthy subjects predict functional alterations in stroke at the level of individual patients. Brain Communications, 6(4), fcae237. 10.1093/braincomms/fcae237

Idesis, S., Favaretto, C., Metcalf, N. V., Griffis, J. C., Shulman, G. L., Corbetta, M., & Deco, G. (2022). Inferring the dynamical effects of stroke lesions through whole-brain modeling. NeuroImage: Clinical, 36, 103233. 10.1016/j.nicl.2022.103233

Kaiser, M., Martin, R., Andras, P., & Young, M. P. (2007). Simulation of robustness against lesions of cortical networks: Simulation of robustness of cortical networks. European Journal of Neuroscience, 25(10), 3185–3192. 10.1111/j.1460-9568.2007.05574.x

Katsari, M., Angelopoulou, G., Laskaris, N., Potagas, C., & Kasselimis, D. (2022). Common Neuroanatomical Substrate of Cholinergic Pathways and Language-Related Brain Regions as an Explanatory Framework for Evaluating the Efficacy of Cholinergic Pharmacotherapy in Post-Stroke Aphasia: A Review. Brain Sciences, 12(10), 1273. 10.3390/brainsci12101273

Kim, Y. K., Yang, E. J., Cho, K., Lim, J. Y., & Paik, N.-J. (2014). Functional Recovery After Ischemic Stroke Is Associated With Reduced GABAergic Inhibition in the Cerebral Cortex: A GABA PET Study. Neurorehabilitation and Neural Repair, 28(6), 576–583. 10.1177/1545968313520411

Ktena, S. I., Schirmer, M. D., Etherton, M. R., Giese, A.-K., Tuozzo, C., Mills, B. B., Rueckert, D., Wu, O., & Rost, N. S. (2019). Brain Connectivity Measures Improve Modeling of Functional Outcome After Acute Ischemic Stroke. Stroke, 50(10), 2761–2767. 10.1161/STROKEAHA.119.025738

Lai, T. W., Zhang, S., & Wang, Y. T. (2014). Excitotoxicity and stroke: Identifying novel targets for neuroprotection. Progress in Neurobiology, 2013 Pangu Meeting on Neurobiology of Stroke and CNS Injury: Progresses and Perspectives of Future, 115, 157–188. 10.1016/j.pneurobio.2013.11.006

Lipkin, B., Tuckute, G., Affourtit, J., Small, H., Mineroff, Z., Kean, H., Jouravlev, O., Rakocevic, L., Pritchett, B., Siegelman, M., Hoeflin, C., Pongos, A., Blank, I. A., Struhl, M. K., Ivanova, A., Shannon, S., Sathe, A., Hoffmann, M., Nieto-Castañón, A., & Fedorenko, E. (2022). Probabilistic atlas for the language network based on precision fMRI data from >800 individuals. Scientific Data, 9(1), 529. 10.1038/s41597-022-01645-3

López-González, A., Panda, R., Ponce-Alvarez, A., Zamora-López, G., Escrichs, A., Martial, C., Thibaut, A., Gosseries, O., Kringelbach, M. L., Annen, J., Laureys, S., & Deco, G. (2021). Loss of consciousness reduces the stability of brain hubs and the heterogeneity of brain dynamics. Communications Biology, 4(1), 1037. 10.1038/s42003-021-02537-9

Lozada, A., Munyao, N., Sallmen, T., Lintunen, M., Leurs, R., Lindsberg, P. J., & Panula, P. (2005). Postischemic regulation of central histamine receptors. Neuroscience, 136(1), 371–379. 10.1016/j.neuroscience.2005.06.079

Markello, R. D., Hansen, J. Y., Liu, Z.-Q., Bazinet, V., Shafiei, G., Suárez, L. E., Blostein, N., Seidlitz, J., Baillet, S., Satterthwaite, T. D., Chakravarty, M. M., Raznahan, A., & Misic, B. (2022). neuromaps: Structural and functional interpretation of brain maps. Nature Methods, 19(11), 1472–1479. 10.1038/s41592-022-01625-w

Mizuki, T., Kobayashi, H., Ueno, S., Nakashima, Y., Kuroiwa, A., & Izumi, F. (1995). Differential Changes in α- and β-Adrenoceptors in the Cerebral Cortex and Hippocampus of the Mongolian Gerbil After Unilateral Brain Ischemia. Stroke, 26(12), 2333–2337. 10.1161/01.STR.26.12.2333

Møller, M., Andersen, G., & Gjedde, A. (2007). Serotonin 5HT1A receptor availability and pathological crying after stroke. Acta Neurologica Scandinavica, 116(2), 83–90. 10.1111/j.1600-0404.2007.00869.x

Mongei Asensio, M. (2025). Modeling the Cortical Landscape of Catecholaminergic Signaling via Spatial Transcriptomics [Universitat Pompeu Fabra]. https://hdl.handle.net/10803/695607

Murphy, T. H., & Corbett, D. (2009). Plasticity during stroke recovery: From synapse to behaviour. Nature Reviews Neuroscience, 10(12), Article 12. 10.1038/nrn2735

Neumann-Haefelin, T., Hagemann, G., & Witte, O. W. (1995). Cellular correlates of neuronal hyperexcitability in the vicinity of photochemically induced cortical infarcts in rats in vitro. Neuroscience Letters, 193(2), 101–104. 10.1016/0304-3940(95)11677-O

Oostenveld, R., Fries, P., Maris, E., & Schoffelen, J.-M. (2011). FieldTrip: Open Source Software for Advanced Analysis of MEG, EEG, and Invasive Electrophysiological Data. Computational Intelligence and Neuroscience, 2011, 1–9. 10.1155/2011/156869

Ponce-Alvarez, A. (2025). Network Mechanisms Underlying the Regional Diversity of Variance and Time Scales of the Brain’s Spontaneous Activity Fluctuations. TheJournal of Neuroscience, 45(10), e1699242024. 10.1523/JNEUROSCI.1699-24.2024

Que, M., Schiene, K., Witte, O. W., & Zilles, K. (1999). Widespread up-regulation of N-methyl-d-aspartate receptors after focal photothrombotic lesion in rat brain.Neuroscience Letters, 273(2), 77–80. 10.1016/S0304-3940(99)00598-4

Que, M., Witte, O. W., Neumann-Haefelin, T., Schiene, K., Schroeter, M., & Zilles, K.(1999). Changes in GABAA and GABAB receptor binding following cortical photothrombosis: A quantitative receptor autoradiographic study. Neuroscience, 93(4), 1233–1240. 10.1016/S0306-4522(99)00197-9

Robb, R. A., & Hanson, D. P. (1991). A software system for interactive and quantitative visualization of multidimensional biomedical images. Australasian Physical & Engineering Sciences in Medicine, 14(1), 9–30.

Sanz Perl, Y., Geli, S., Pérez-Ordoyo, E., Zonca, L., Idesis, S., Vohryzek, J., Jirsa, V. K., Kringelbach, M. L., Tagliazucchi, E., & Deco, G. (2023). Whole-brain modelling of low-dimensional manifold modes reveals organising principle of brain dynamics. 10.1101/2023.11.20.567824

Sanz Perl, Y., Zamora-Lopez, G., Montbrió, E., Monge-Asensio, M., Vohryzek, J., Fittipaldi, S., Campo, C. G., Moguilner, S., Ibañez, A., Tagliazucchi, E., Yeo, B. T. T., Kringelbach, M. L., & Deco, G. (2023). The impact of regional heterogeneity in whole-brain dynamics in the presence of oscillations. Network Neuroscience, 7(2), 632–660. 10.1162/netn_a_00299

Sarasso, S., D’Ambrosio, S., Fecchio, M., Casarotto, S., Viganò, A., Landi, C., Mattavelli, G., Gosseries, O., Quarenghi, M., Laureys, S., Devalle, G., Rosanova, M., & Massimini, M. (2020). Local sleep-like cortical reactivity in the awake brain after focal injury. Brain: A Journal of Neurology, 143(12), 3672–3684. 10.1093/brain/awaa338

Schaefer, A., Kong, R., Gordon, E. M., Laumann, T. O., Zuo, X.-N., Holmes, A. J., Eickhoff, S. B., & Yeo, B. T. T. (2018). Local-Global Parcellation of the Human Cerebral Cortex from Intrinsic Functional Connectivity MRI. Cerebral Cortex (New York, N.Y.: 1991), 28(9), 3095–3114. 10.1093/cercor/bhx179

Schiene, K., Bruehl, C., Zilles, K., Qu, M., Hagemann, G., Kraemer, M., & Witte, O. W. (1996). Neuronal Hyperexcitability and Reduction of GABAA-Receptor Expression in the Surround of Cerebral Photothrombosis. Journal of Cerebral Blood Flow & Metabolism, 16(5), 906–914. 10.1097/00004647-199609000-00014

Schirmer, M. D., Ktena, S. I., Nardin, M. J., Donahue, K. L., Giese, A.-K., Etherton, M. R., Wu, O., & Rost, N. S. (2019). Rich-Club Organization: An Important Determinant of Functional Outcome After Acute Ischemic Stroke. Frontiers in Neurology, 10, 956. 10.3389/fneur.2019.00956

Schulte, J., Senden, M., Deco, G., Kobeleva, X., & Zamora-López, G. (2025). The global communication pathways of the human brain transcend the cortical-subcortical-cerebellar division (Version 3). arXiv. 10.48550/ARXIV.2505.22893

Sporns, O., Honey, C. J., & Kötter, R. (2007). Identification and Classification of Hubs in Brain Networks. PLoS ONE, 2(10), e1049. 10.1371/journal.pone.0001049

Stam, C. J. (2024). Hub overload and failure as a final common pathway in neurological brain network disorders. Network Neuroscience, 8(1), 1–23. 10.1162/netn_a_00339

Stephan, K. E., Weiskopf, N., Drysdale, P. M., Robinson, P. A., & Friston, K. J. (2007).Comparing hemodynamic models with DCM. NeuroImage, 38(3), 387–401. 10.1016/j.neuroimage.2007.07.040

Tzourio-Mazoyer, N., Landeau, B., Papathanassiou, D., Crivello, F., Etard, O., Delcroix, N., Mazoyer, B., & Joliot, M. (2002). Automated anatomical labeling of activations inSPM using a macroscopic anatomical parcellation of the MNI MRI single-subject brain. NeuroImage, 15(1), 273–289. 10.1006/nimg.2001.0978

Van Den Heuvel, M. P., & Sporns, O. (2013). Network hubs in the human brain. Trends in Cognitive Sciences, 17(12), 683–696. 10.1016/j.tics.2013.09.012

Véra, P., Zilbovicius, M., Chabriat, H., Amarenco, P., Kerdraon, J., Ménard, J. F., Amarenco, G., Bousser, M. G., Rancurel, G., & Samson, Y. (1996). Post-stroke changes in cortical 5-HT2 serotonergic receptors. Journal of Nuclear Medicine: Official Publication, Society of Nuclear Medicine, 37(12), 1976–1981.

Wagner, H. H., & Dray, S. (2015). Generating spatially constrained null models for irregularly spaced data using M oran spectral randomization methods. Methods in Ecology and Evolution, 6(10), 1169–1178. 10.1111/2041-210X.12407

Wang, L. E., Fink, G. R., Diekhoff, S., Rehme, A. K., Eickhoff, S. B., & Grefkes, C. (2011). Noradrenergic enhancement improves motor network connectivity in stroke patients. Annals of Neurology, 69(2), 375–388. 10.1002/ana.22237

Wang, L., Xu, X., Kai Lau, K., Li, L. S. W., Kwun Wong, Y., Yau, C., Mak, H. K. F., & Hui,E. S. (2021). Relation between rich-club organization versus brain functions and functional recovery after acute ischemic stroke. Brain Research, 1763, 147441. 10.1016/j.brainres.2021.147441

Ward, N. S. (2017). Restoring brain function after stroke—Bridging the gap between animals and humans. Nature Reviews Neurology, 13(4), 244–255. 10.1038/nrneurol.2017.34

Wong, K.-F., & Wang, X.-J. (2006). A Recurrent Network Mechanism of Time Integration in Perceptual Decisions. The Journal of Neuroscience, 26(4), 1314–1328. 10.1523/JNEUROSCI.3733-05.2006

Yang, L., Wang, Y., & Chen, Z. (2022). Central histaminergic signalling, neural excitability and epilepsy. British Journal of Pharmacology, 179(1), 3–22. 10.1111/bph.15692

Yeh, F.-C., Panesar, S., Fernandes, D., Meola, A., Yoshino, M., Fernandez-Miranda, J. C., Vettel, J. M., & Verstynen, T. (2018). Population-averaged atlas of the macroscale human structural connectome and its network topology. NeuroImage, 178, 57–68. 10.1016/j.neuroimage.2018.05.027

Yeh, F.-C., & Tseng, W.-Y. I. (2011). NTU-90: A high angular resolution brain atlas constructed by q-space diffeomorphic reconstruction. NeuroImage, 58(1), 91–99. 10.1016/j.neuroimage.2011.06.021

Zahrai, A., Vahid-Ansari, F., Daigle, M., & Albert, P. R. (2020). Fluoxetine-induced recovery of serotonin and norepinephrine projections in a mouse model of post-stroke depression. Translational Psychiatry, 10(1), 334. 10.1038/s41398-020-01008-9

Zamora-López, G., Zhou, C., & Kurths, J. (2009). Graph analysis of cortical networks reveals complex anatomical communication substrate. Chaos: An Interdisciplinary Journal of Nonlinear Science, 19(1), 015117. 10.1063/1.3089559

Zamora-López, G., Zhou, C., & Kurths, J. (2010). Cortical hubs form a module for multisensory integration on top of the hierarchy of cortical networks. Frontiers in Neuroinformatics. 10.3389/neuro.11.001.2010

Zamora-López, G., Zhou, C., & Kurths, J. (2011). Exploring Brain Function from Anatomical Connectivity. Frontiers in Neuroscience, 5, 83. 10.3389/fnins.2011.00083

