## Supplementary Materials Whole-brain receptor markers and hub excitabilities predict acute and long-term outcomes in stroke for "Whole-brain receptor markers and hub excitabilities determine acute and long-term outcomes in stroke"

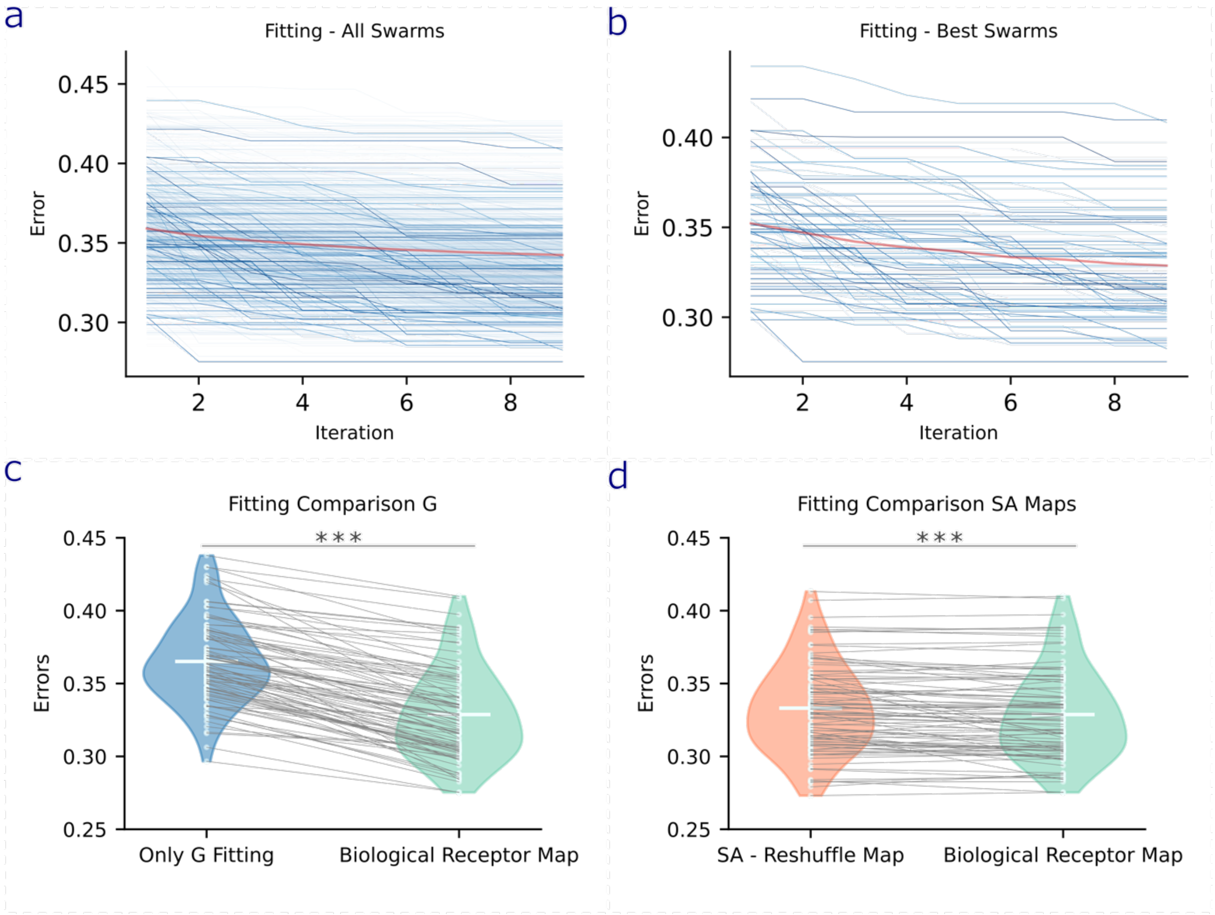


**S1 – Whole-brain model fitting results.** **a |** For each subject, we applied a multi–particle swarm optimization (PSO) procedure with 10 swarms each containing 60 particles and a maximum of 9 iterations. **b |** The best-performing swarm for each patient was selected as the final model solution. **c |** Model fits obtained through PSO yielded significantly lower errors compared to those derived from the initial global coupling factor optimization (*$p \leq 0.05$, **$p \leq 0.01$, ***$p \leq 0.001$). **d |** We further compared these fits to those obtained using spatially autocorrelated surrogate receptor maps and observed significantly better performance with the empirical biological receptor maps, underscoring the topographical relevance of the true receptor distributions.

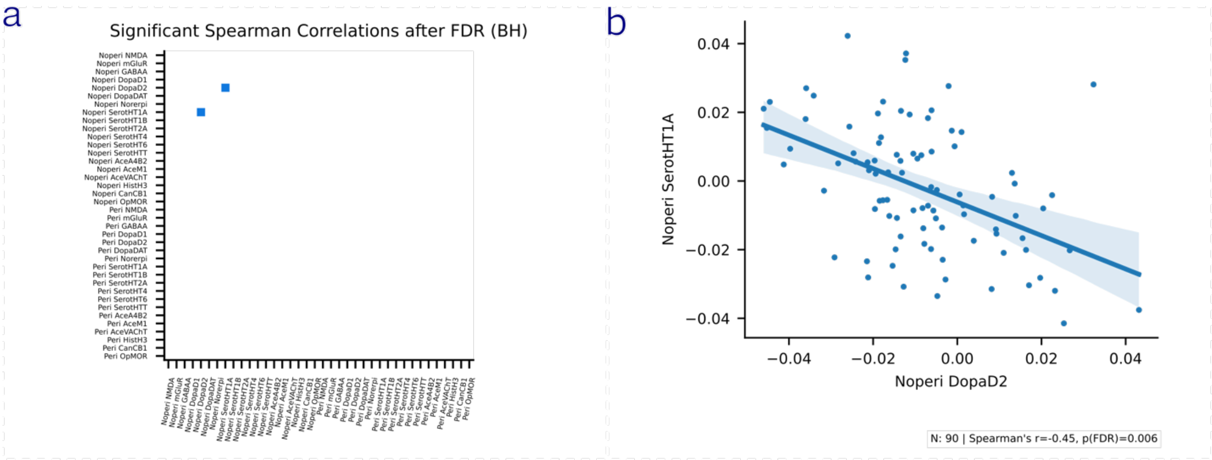


**Figure S2 – Significant correlations between receptor scalings. a & b |** After correcting for multiple comparisons, we found a significant anticorrelation between non-perilesional DopaD2 and non-perilesional serotonin HT1A receptor scalings.

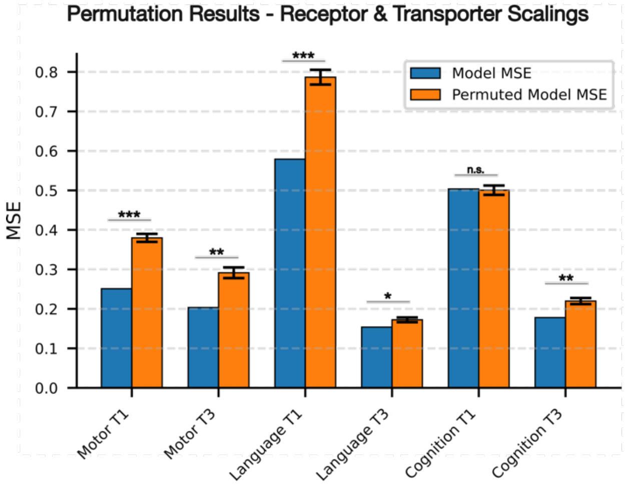


**Figure S3 – Ridge Regression permutation results (LOOCV).** Synaptic receptor and transporter scalings were used to explain acute (T1) and long-term (T3) motor, language, and cognitive outcomes. Observed mean squared errors (MSE; blue) are shown alongside the null distribution obtained by permuting the dependent variable across 10,000 iterations (orange; SD indicated). After correction for multiple comparisons, all models except Cognition T1 reached significance (*$p \leq 0.05$, **$p \leq 0.01$, ***$p \leq 0.001$).

| Model | N | R2 | R2 Perm. Mean | R2 Perm. Std. | MSE | MSE Perm. Mean | MSE Perm. Std. | Model P-val. | Model P-val.(FDR) |
|---|---|---|---|---|---|---|---|---|---|
| **Motor T1** | 78 | 0.33 | -0.0144 | 0.0271 | 0.251 | 0.38 | 0.0101 | 0.0002 | 0.0006 |

| | | | | | | | | | |
|---|---|---|---|---|---|---|---|---|---|
| **Motor T3** | 47 | 0.286 | -0.0236 | 0.0477 | 0.203 | 0.291 | 0.0136 | 0.0033 | 0.0066 |
| **Language T1** | 84 | 0.254 | -0.0138 | 0.0241 | 0.579 | 0.787 | 0.0187 | 0.0002 | 0.0006 |
| **Language T3** | 54 | 0.0866 | -0.0235 | 0.0357 | 0.154 | 0.172 | 0.006 | 0.0202 | 0.0242 |
| **Cognition T1** | 86 | -0.0195 | -0.0132 | 0.0239 | 0.504 | 0.5 | 0.0118 | 0.321 | 0.321 |
| **Cognition T3** | 55 | 0.173 | -0.0213 | 0.0362 | 0.178 | 0.22 | 0.00779 | 0.0051 | 0.00765 |

**Table ST1 - Model performance of the ridge regressions (LOOCV) and corresponding permutation results using synaptic receptor and transporter scalings as predictors together with covariates.** Model p-values reflect the probability of observing the model's mean squared error (MSE) under the permuted null distribution. All p-values were also corrected for multiple comparisons.

| Predictor | Coeff. | Permutation Coeff. Mean | Permutation Coeff. Std. | P-val. | P-val. (FDR) |
|---|---|---|---|---|---|
| **Peri NMDA** | 0.029 | 0.000 | 0.010 | 0.030 | 0.047 |
| **Peri mGluR** | -0.035 | 0.000 | 0.010 | 0.018 | 0.030 |
| **Peri DopaD2** | 0.048 | 0.000 | 0.010 | 0.008 | 0.019 |
| **Peri DopaDAT** | -0.073 | 0.000 | 0.010 | 0.002 | 0.009 |
| **Peri NET** | 0.092 | -0.000 | 0.010 | 0.001 | 0.006 |
| **Peri SerotHT1B** | -0.054 | 0.000 | 0.011 | 0.007 | 0.017 |
| **Peri SerotHT4** | 0.043 | 0.000 | 0.011 | 0.014 | 0.024 |
| **Peri SerotHT6** | -0.044 | 0.000 | 0.011 | 0.011 | 0.022 |
| **Peri SerotHTT** | 0.054 | 0.000 | 0.010 | 0.005 | 0.014 |
| **Peri AceA4B2** | -0.067 | 0.000 | 0.010 | 0.002 | 0.009 |
| **Peri AceM1** | 0.054 | 0.000 | 0.011 | 0.006 | 0.016 |
| **Peri HistH3** | 0.092 | -0.000 | 0.010 | 0.001 | 0.009 |
| **Peri CanCB1** | 0.042 | -0.000 | 0.010 | 0.012 | 0.022 |
| **Peri OpMOR** | -0.061 | -0.000 | 0.010 | 0.002 | 0.009 |
| **NoPeri NMDA** | -0.071 | 0.000 | 0.010 | 0.002 | 0.009 |
| **NoPeri GABAA** | -0.115 | 0.000 | 0.010 | 0.000 | 0.004 |
| **NoPeri DopaD1** | 0.050 | -0.000 | 0.010 | 0.007 | 0.017 |
| **NoPeri DopaDAT** | -0.039 | -0.000 | 0.010 | 0.013 | 0.023 |
| **NoPeri SerotHT2A** | -0.040 | 0.000 | 0.010 | 0.012 | 0.022 |
| **NoPeri SerotHT6** | 0.054 | -0.000 | 0.010 | 0.005 | 0.014 |
| **NoPeri SerotHTT** | -0.029 | -0.000 | 0.010 | 0.028 | 0.046 |
| **NoPeri AceVAChT** | 0.069 | -0.000 | 0.010 | 0.002 | 0.009 |
| **NoPeri HistH3** | 0.063 | 0.000 | 0.010 | 0.003 | 0.010 |
| **NoPeri CanCB1** | 0.090 | -0.000 | 0.010 | 0.001 | 0.006 |
| **NoPeri OpMOR** | -0.060 | -0.000 | 0.010 | 0.003 | 0.011 |
| **Expert localization_cerebellar** | 0.081 | 0.000 | 0.011 | 0.002 | 0.009 |
| **Expert localization_cortical** | 0.111 | -0.000 | 0.010 | 0.000 | 0.004 |
| **Expert localization_other** | -0.043 | 0.000 | 0.010 | 0.009 | 0.020 |
| **PCA A Nodal Damage 0** | -0.176 | -0.000 | 0.010 | 0.000 | 0.004 |
| **PCA A Nodal Damage 1** | -0.038 | 0.000 | 0.008 | 0.008 | 0.019 |
| **PCA A Nodal Damage 2** | 0.114 | 0.000 | 0.010 | 0.000 | 0.004 |
| **PCA A Nodal Damage 4** | -0.041 | 0.000 | 0.009 | 0.010 | 0.020 |

**Table ST2 - Significant model predictors of the synaptic receptor and transporter ridge regression analysis predicting acute motor performance.**

| Predictor | Coeff. | Permutation Coeff. Mean | Permutation Coeff. Std. | P-val. | P-val. (FDR) |
|---|---|---|---|---|---|
| Peri NMDA | 0.092 | 0.000 | 0.014 | 0.002 | 0.020 |
| Peri DopaD1 | -0.071 | -0.000 | 0.014 | 0.006 | 0.025 |
| Peri SerotHT1B | -0.117 | -0.000 | 0.014 | 0.001 | 0.015 |
| Peri SerotHT4 | 0.076 | -0.000 | 0.014 | 0.005 | 0.025 |
| Peri SerotHT6 | 0.062 | 0.000 | 0.014 | 0.010 | 0.034 |
| Peri SerotHTT | 0.072 | 0.000 | 0.013 | 0.005 | 0.025 |
| Peri AceVAChT | -0.101 | 0.000 | 0.014 | 0.001 | 0.015 |
| NoPeri NMDA | -0.116 | 0.000 | 0.014 | 0.001 | 0.015 |
| NoPeri DopaDAT | -0.073 | 0.000 | 0.013 | 0.005 | 0.025 |
| NoPeri SerotHT2A | -0.047 | 0.000 | 0.013 | 0.014 | 0.047 |
| NoPeri AceM1 | 0.065 | -0.000 | 0.013 | 0.006 | 0.025 |
| NoPeri AceVAChT | 0.079 | 0.000 | 0.014 | 0.005 | 0.025 |
| NoPeri CanCB1 | 0.048 | 0.000 | 0.013 | 0.016 | 0.048 |
| Expert localization_cerebellar | 0.044 | -0.000 | 0.014 | 0.018 | 0.048 |
| Expert localization_cortico_subcortical | -0.059 | 0.000 | 0.012 | 0.006 | 0.025 |
| Expert localization_white matter only | 0.047 | -0.000 | 0.015 | 0.016 | 0.048 |
| PCA A Nodal Damage 0 | -0.266 | 0.000 | 0.012 | 0.000 | 0.002 |
| PCA A Nodal Damage 1 | 0.273 | 0.000 | 0.014 | 0.000 | 0.002 |
| PCA A Nodal Damage 3 | 0.042 | 0.000 | 0.012 | 0.017 | 0.048 |

**Table ST3 - Significant model predictors of the synaptic receptor and transporter ridge regression analysis predicting acute language performance.**

| Predictor | Beta (standardized) | Std. Error | z | P-val. | [0.025 | 0.975] | P (FDR-corrected) | Significant |
|---|---|---|---|---|---|---|---|---|
| const | -0.029 | 0.157 | -0.182 | 0.856 | -0.337 | 0.279 | - | - |
| A LH Default Temp 3 | 0.324 | 0.11 | 2.946 | 0.003 | 0.108 | 0.539 | 0.045 | True |
| Nodal Damage LH Default Temp 3 | -0.089 | 0.278 | -0.32 | 0.749 | -0.633 | 0.455 | 0.874 | False |
| A LH SomMot 11 | -0.166 | 0.104 | -1.594 | 0.111 | -0.37 | 0.038 | 0.264 | False |
| Nodal Damage LH SomMot 11 | -0.038 | 0.162 | -0.237 | 0.812 | -0.355 | 0.278 | 0.875 | False |
| A RH DorsAttn Post 1 | -0.327 | 0.137 | -2.389 | 0.017 | -0.595 | -0.059 | 0.118 | False |
| Nodal Damage RH DorsAttn Post 1 | 0.437 | 0.29 | 1.507 | 0.132 | -0.131 | 1.005 | 0.264 | False |
| A RH Cont Par 3 | 0.01 | 0.165 | 0.06 | 0.952 | -0.313 | 0.333 | 0.952 | False |
| Nodal Damage RH Cont Par 3 | -0.149 | 0.254 | -0.587 | 0.557 | -0.648 | 0.349 | 0.78 | False |

| | | | | | | | | |
|---|---|---|---|---|---|---|---|---|
| **A RH DorsAttn Post 8** | 0.235 | 0.143 | 1.641 | 0.101 | -0.046 | 0.516 | 0.264 | False |
| **Nodal Damage RH DorsAttn Post 8** | 0.263 | 0.277 | 0.951 | 0.342 | -0.28 | 0.806 | 0.598 | False |
| **PCA_nodal_0** | -0.802 | 0.416 | -1.929 | 0.054 | -1.618 | 0.013 | 0.251 | False |
| **PCA_nodal_1** | 0.376 | 0.241 | 1.562 | 0.118 | -0.096 | 0.848 | 0.264 | False |
| **PCA_nodal_2** | -0.22 | 0.311 | -0.706 | 0.48 | -0.829 | 0.39 | 0.747 | False |
| **PCA_nodal_3** | 0.082 | 0.18 | 0.458 | 0.647 | -0.27 | 0.435 | 0.824 | False |
| **Lesion Etiology_Ischemic** | 0.183 | 0.178 | 1.025 | 0.305 | -0.167 | 0.533 | - | - |

**Table ST4 - Multivariate linear regression analysis of acute language performance (T1).** Standardized regression coefficients (*β*), heteroscedasticity-robust standard errors (HC3), and corresponding z-statistics are reported for all model predictors (*N* = 90). P values for the variables of interest were adjusted for multiple comparisons using the Benjamini–Hochberg False Discovery Rate (FDR) procedure. Significance was defined as *P(FDR)* < 0.05. Confidence intervals (95% CI) are provided for the standardized betas. Predictors include hub excitability (A), hub nodal damage, and principal components of the nodal damage of all non-hubs (PCA 0–3).

| Predictor | Beta (standardized) | Std. Error | t | P-val. | [0.025 | 0.975] | P (FDR-corrected) | Significant |
|---|---|---|---|---|---|---|---|---|
| **const** | -0.154 | 0.19 | -0.809 | 0.421 | -0.533 | 0.225 | - | - |
| **A LH Default Temp 3** | 0.363 | 0.099 | 3.681 | 0.0 | 0.167 | 0.56 | 0.002 | True |
| **Nodal Damage LH Default Temp 3** | 0.077 | 0.173 | 0.442 | 0.66 | -0.269 | 0.422 | 0.66 | False |
| **A LH SomMot 11** | -0.394 | 0.102 | -3.86 | 0.0 | -0.598 | -0.191 | 0.002 | True |
| **Nodal Damage LH SomMot 11** | 0.216 | 0.098 | 2.2 | 0.031 | 0.02 | 0.412 | 0.066 | False |
| **A * Nodal Damage LH SomMot 11** | 0.2 | 0.064 | 3.121 | 0.003 | 0.072 | 0.327 | 0.008 | True |
| **A RH DorsAttn Post 1** | -0.245 | 0.103 | -2.385 | 0.02 | -0.45 | -0.04 | 0.049 | True |
| **Nodal Damage RH DorsAttn Post 1** | 0.528 | 0.144 | 3.675 | 0.0 | 0.242 | 0.814 | 0.002 | True |
| **A RH Cont Par 3** | 0.109 | 0.132 | 0.829 | 0.41 | -0.153 | 0.372 | 0.512 | False |
| **Nodal Damage RH Cont Par 3** | -0.165 | 0.167 | -0.989 | 0.326 | -0.498 | 0.168 | 0.445 | False |
| **A RH DorsAttn Post 8** | 0.217 | 0.123 | 1.763 | 0.082 | -0.028 | 0.462 | 0.137 | False |
| **Nodal Damage RH DorsAttn Post 8** | 0.246 | 0.164 | 1.498 | 0.138 | -0.081 | 0.574 | 0.207 | False |
| **PCA_nodal_0** | -0.873 | 0.243 | -3.595 | 0.001 | -1.357 | -0.389 | 0.002 | True |
| **PCA_nodal_1** | 0.105 | 0.162 | 0.65 | 0.518 | -0.217 | 0.427 | 0.555 | False |

| | | | | | | | | |
|---|---|---|---|---|---|---|---|---|
| **PCA_nodal_2** | -0.281 | 0.135 | -2.079 | 0.041 | -0.55 | -0.012 | 0.077 | False |
| **PCA_nodal_3** | -0.057 | 0.08 | -0.718 | 0.475 | -0.216 | 0.102 | 0.548 | False |
| **Lesion Etiology_Ischemic** | 0.117 | 0.205 | 0.571 | 0.57 | -0.291 | 0.525 | - | - |

**Table ST5 - Multivariate linear regression analysis of acute cognitive performance (T1).** Standardized regression coefficients ($\beta$), standard errors, and t-statistics are reported ($N = 92$). P-values for variables of interest were FDR-corrected (Benjamini–Hochberg); significance was defined as *P(FDR)* < 0.05. Model predictors include hub excitability (A), nodal damage, excitability × damage interactions, and non-hub nodal damage pc scores (PCA 0–3). *Note: The significant effect of Nodal Damage RH DorsAttn Post 1 was found to be outlier-driven and became non-significant upon their removal.

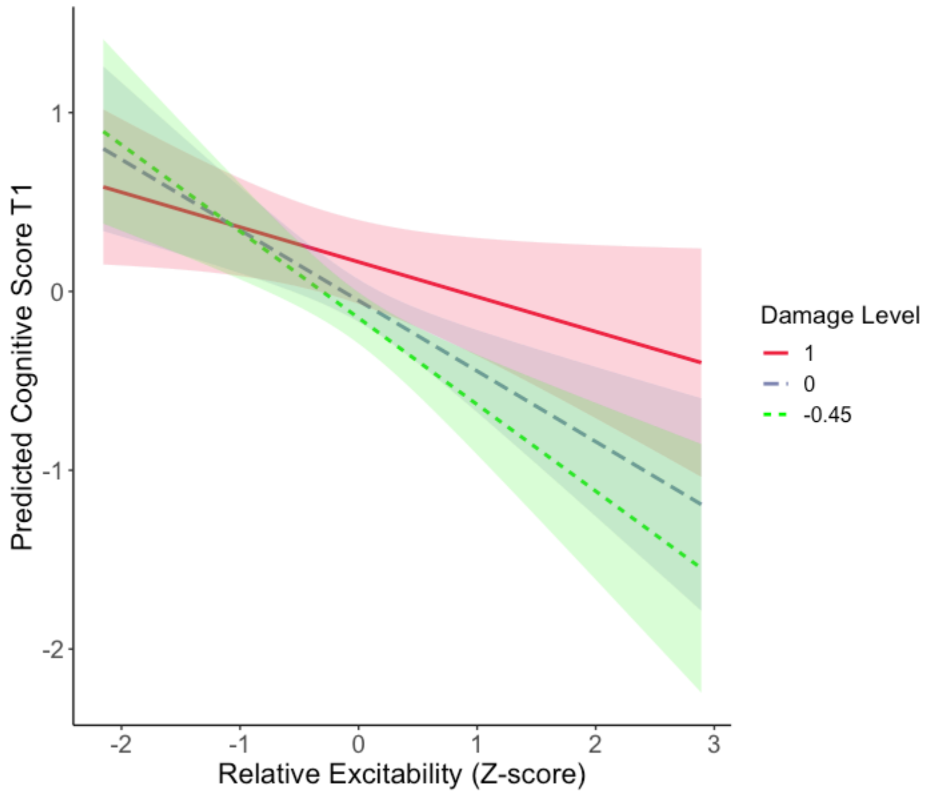


**Figure S4 - Cognition T1: Interaction in LH SomMot 11.** Simple slope plots demonstrate that higher excitability significantly predicts lower acute cognitive scores in patients with no nodal damage (-0.45 *SD*, green; $\beta = -0.48$, $t = -4.14$, $p < 0.001$) and average damage (blue; $\beta = -0.39$, $t = -3.86$, $p < 0.001$). This negative association is substantially attenuated in patients with high structural damage (+1 *SD*, red; $\beta = -0.19$, $t = -2.03$, $p = 0.05$), suggesting that the detrimental effect of excitability on acute function is most pronounced in structurally intact nodes.

| Predictor | Beta (standardized) | Std. Error | t | P-val. | [0.025 | 0.975] | P (FDR-corrected) | Significant |
|---|---|---|---|---|---|---|---|---|
| **const** | 0.345 | 0.13 | 2.653 | 0.011 | 0.082 | 0.608 | - | - |
| **Language_t1** | 0.309 | 0.048 | 6.452 | 0.0 | 0.212 | 0.406 | 0.0 | True |
| **A LH Default Temp 3** | 0.007 | 0.055 | 0.125 | 0.901 | -0.103 | 0.117 | 0.951 | False |
| **Nodal Damage LH Default Temp 3** | 0.067 | 0.106 | 0.628 | 0.533 | -0.148 | 0.281 | 0.8 | False |
| **A LH SomMot 11** | 0.012 | 0.052 | 0.225 | 0.823 | -0.093 | 0.116 | 0.951 | False |
| **Nodal Damage LH SomMot 11** | -0.015 | 0.055 | -0.279 | 0.782 | -0.126 | 0.095 | 0.951 | False |
| **A RH DorsAttn Post 1** | 0.162 | 0.058 | 2.818 | 0.007 | 0.046 | 0.278 | 0.037 | True |
| **Nodal Damage RH DorsAttn Post 1** | -0.368 | 0.143 | -2.573 | 0.014 | -0.657 | -0.079 | 0.052 | False |
| **A * Nodal Damage RH DorsAttn Post 1** | 0.197 | 0.061 | 3.237 | 0.002 | 0.074 | 0.32 | 0.018 | True |
| **A RH Cont Par 3** | -0.064 | 0.069 | -0.928 | 0.359 | -0.204 | 0.075 | 0.769 | False |
| **Nodal Damage RH Cont Par 3** | 0.207 | 0.114 | 1.818 | 0.076 | -0.023 | 0.438 | 0.229 | False |
| **A RH DorsAttn Post 8** | -0.004 | 0.068 | -0.062 | 0.951 | -0.142 | 0.134 | 0.951 | False |
| **Nodal Damage RH DorsAttn Post 8** | 0.061 | 0.09 | 0.682 | 0.499 | -0.12 | 0.243 | 0.8 | False |
| **PCA_nodal_1** | -0.163 | 0.108 | -1.511 | 0.138 | -0.38 | 0.055 | 0.346 | False |
| **PCA_nodal_2** | -0.05 | 0.066 | -0.759 | 0.452 | -0.183 | 0.083 | 0.8 | False |
| **PCA_nodal_3** | -0.008 | 0.059 | -0.128 | 0.899 | -0.127 | 0.111 | 0.951 | False |
| **Lesion Etiology_Ischemic** | 0.037 | 0.134 | 0.277 | 0.783 | -0.234 | 0.308 | - | - |

**Table ST6 - Multivariate linear regression analysis of long-term language performance (T3).** Standardized regression coefficients ($\beta$), standard errors, and t-statistics are reported ($N = 58$). P-values for variables of interest were FDR-corrected (Benjamini–Hochberg); significance was defined as *P(FDR)* < 0.05. Model predictors include initial language performance, hub excitability (A), nodal damage, excitability × damage interactions, and non-hub nodal damage pc scores (PCA 1–3). PCA 0 scores were dropped due to high multicollinearity.

| Parameter | Value |
|---|---|
| $\kappa$ | 1 / 0.65 |
| $\gamma$ | 1 / 0.41 |
| $\tau$ | 0.98 |
| $\alpha$ | 0.32 |
| $k_1$ | $4.3\mathscr{V}_0 E_0 TE$ |
| $k_1$ | $r_0 E_0 TE$ |
| $k_1$ | 1 |
| $\mathscr{V}_0$ | 40.3 |
| $E_0$ | 0.4 |
| $TE$ | 0.04 |
| $r_0$ | 25 |

**Table ST7 | Biophysical parameters of the Hemodynamic Model.**
